# Systematic investigation reveals extensive Epstein-Barr virus transcriptional regulation of the human genome

**DOI:** 10.1101/2025.07.18.665561

**Authors:** Phillip J. Dexheimer, Matthew R. Hass, Lee E. Edsall, Arame A. Diouf, Sydney H. Jones, Omer Donmez, Cailing Yin, Katelyn A. Dunn, Carmy Forney, Hayley K. Hesse, Andrew VonHandorf, Xiaoting Chen, Sreeja Parameswaran, Olivia E. Gittens, Kenyatta C. M. F. Viel, Rozita Razavi, Benjamin E. Gewurz, Bo Zhao, Lucinda P. Lawson, Timothy R. Hughes, Leah C. Kottyan, Matthew T. Weirauch

## Abstract

We systematically investigate interactions between Epstein-Barr virus (EBV) transcriptional regulators (vTRs) and the human genome. Starting with 16 known and candidate vTRs, we identify nine whose introduction into human cells results in substantial alterations to host gene expression. Genome-scale determination of vTR genomic binding events and alterations to chromatin accessibility reveals a detailed map of EBV’s functional interactions with the human genome, including >100,000 vTR binding events impacting almost a quarter of all human genes. BMRF1 emerges as a potent regulator, impacting >7,000 genes and altering >37,000 chromatin regions. Our results provide new evidence that EBV RTA interacts with and stabilizes the binding of human RBPJ. Network analysis reveals that many human genes are targeted by multiple EBV vTRs, highlighting the vast coordinated impact of EBV on human gene expression. This study provides a valuable, extensive resource for examining EBV-induced alterations to human gene regulation, with data available on multiple platforms.

## Introduction

Epstein-Barr virus (EBV) is a near-ubiquitous gammaherpesvirus with an established, causative role in mononucleosis (1), Burkitt lymphoma (2), Hodgkin lymphoma (3), and nasopharyngeal carcinoma (4). Compelling emerging evidence also implicates EBV in several autoimmune diseases, including multiple sclerosis (5), systemic lupus erythematosus (6, 7), rheumatoid arthritis (8–10), and inflammatory bowel disease (11). Multiple EBV-based mechanisms are known to contribute to disease processes, including transformation of B cells into malignant cells in cancers (12–14), molecular mimicry between EBV-encoded EBNA1 and GlialCAM in multiple sclerosis (15), and altered regulation of human genes by EBV-encoded EBNA2 in multiple autoimmune diseases (16–19). Despite these advances, it is likely that many additional mechanisms contribute to EBV-based disease pathogenesis which have not yet been identified.

EBV survives by manipulating host processes on multiple levels (20–22). EBV encodes 16 known or suspected viral transcriptional regulators (vTRs), including both direct and indirect DNA binding proteins). EBV has the second most vTRs of the 419 known human viruses (23). For some of these proteins, such as EBNA2 and ZTA (also known as Z, ZEBRA, or BZLF1), their viral and human genome targets have been characterized by the community over decades of studies (16–19, 24–41). Others, such as RTA (also known as R or BRLF1) and BMRF1, remain more elusive in the context of their interactions with the human genome. Even for the well-studied proteins, differences in assays, protocols, data processing, and cell types make direct comparison of EBV vTR mechanisms a challenge. Further, a small subset of EBV vTRs currently have published ChIP-seq data mapped to the human genome in any context (16–18, 37, 42–54). To fully understand the complex layers of gene regulation used by EBV to manipulate human gene expression levels, a uniform systems-level approach is therefore needed.

Here, we employed three functional genomics assays aimed at systematically characterizing the gene regulatory targets and mechanisms of known and suspected EBV vTRs (23). We first examined the transcriptional role of each vTR by transiently expressing each vTR one at a time in human cells. After confirming human gene expression impact (RNA-seq), we assessed the effects of stable vTR expression on human gene expression (RNA-seq), genome binding (ChIP-seq) and chromatin accessibility (ATAC-seq). Altogether, our work encompasses 106 transient RNA-seq experiments, 33 stable RNA-seq experiments, 30 ATAC-seq experiments, and 29 ChIP-seq experiments. We also performed nine ChIP-seq experiments for the human-encoded transcription factor RBPJ due to its known interaction with EBNA2 and its critical role in EBV biology (18). Through a variety of approaches, we confirm that our system produces high quality data, capable of capturing known EBV biology.

The result is an unprecedented dataset that reveals a staggering impact of EBV-encoded vTRs on the human genome, with >15,000 significantly altered human gene expression levels, >64,000 unique vTR-occupied regions of the human genome, and >109,000 regions with significantly altered chromatin accessibility. Based on comprehensive human Transcription Factor (TF) motif enrichment analysis, we discover that the regions directly bound and opened by each EBV vTR are strongly enriched for AP-1 family motifs, whereas the regions directly bound and closed by eight of the nine vTRs are strongly enriched for CTCF motifs. We predict and confirm that, similar to EBNA2, RTA interacts with the human-encoded RBPJ TF, stabilizing its binding at thousands of human genomic loci. Strikingly, the relatively understudied EBV-encoded BMRF1 protein alters the expression of >7,000 human genes, occupies >12,000 regions of the human genome, and alters chromatin accessibility at >37,000 regions. BMRF1 binding has important implications for human disease; for example, regions occupied by BMRF1 are highly enriched for genetic variants associated with rheumatoid arthritis (RA). Systems-level modeling of vTR interactions revealed a complex landscape where many human genes are targeted by multiple vTRs, including *MYC*, *BCL11B*, and *STAT1*.

Collectively, these data comprise a valuable resource for the systematic study of the impact of EBV vTRs on human gene regulatory mechanisms. The data are freely available on multiple platforms, facilitating future studies aimed at understanding EBV vTR binding events, impacts on chromatin accessibility, alterations of human gene expression, and connections to EBV-implicated human diseases.

## Results

### An experimental system to systematically investigate EBV vTR-induced impacts on human gene regulation

We initially sought to identify the role of vTRs encoded by EBV on human transcription. To this end, we focused on eight previously described vTRs (EBNA1, EBNA2, EBNA3A, EBNA3B, EBNA3C, EBNALP, RTA, and ZTA) and eight less characterized vTRs (BALF2, BCRF1, BDLF3, BDLF4, BFRF2, BGLF3, BMRF1, and BVLF1) that were systematically identified in our catalogue of vTRs from all human viruses (23). We transiently expressed each EBV vTR and measured the impact on human gene expression relative to a control cell line that only expressed GFP (**Figure 1A**). Of the eight less characterized EBV vTRs, we selected BMRF1 for further study based on its robust and non-redundant impact on gene expression (**Supplemental Figure 1**). Notably, BMRF1 demonstrated the most robust impact on gene expression in the transient system (**Supplemental Figure 1 panels A and B**), with strong separation from controls and the other less characterized vTRs in a principal component analysis (PCA) (**Supplemental Figure 1 panel C**).

**Figure 1.**
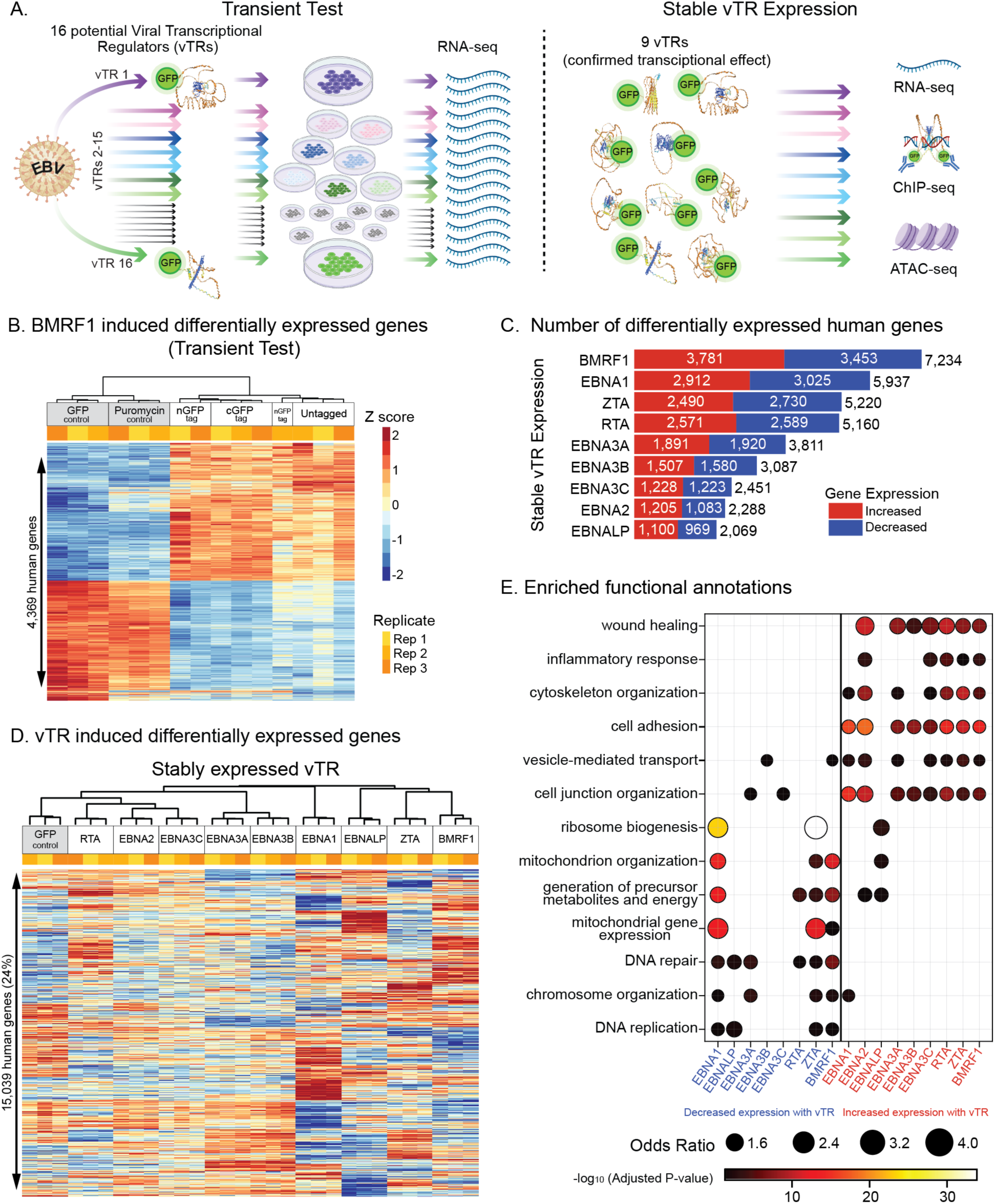
EBV vTRs induce extensive non-redundant changes to human gene expression. **A.** Experimental system. **B.** Differential human gene expression in the transient test of one representative vTR (BMRF1). BMRF1-induced differentially expressed human genes are shown for experimental triplicates. Values indicate normalized gene expression levels that have been Z-transformed across rows (conditions). Columns and rows were clustered using Ward’s method. **C.** Number of differentially expressed human genes in cells stably expressing each vTR. **D.** Quantification of vTR induced human gene expression changes in cells stably expressing each vTR. All human genes with statistically reproducible differential expression induced by any vTR are shown. Values indicate normalized gene expression levels that have been Z-transformed across rows (conditions). Columns and rows were clustered using Ward’s method. **E.** Functional annotations enriched among up and down regulated genes across cell lines stably expressing each vTR. Enriched pathways were identified using ToppGene (see Methods). Up to the top five most significant GO Slim Biological Process categories for each vTR are included. Panel has been curated to reduce redundant pathways (see **Supplemental Figure 7** for full results).

We next performed a series of experiments aimed at optimizing our reagents for downstream use. In this study, we use GFP as an epitope tag to allow us to perform chromatin immunoprecipitation to measure genome-wide binding (**Figure 1A**). We performed transient transfection tests in which GFP was tagged to each vTR on either the C- or the N-terminus. Using RNA sequencing (RNA-seq), we measured gene expression with and without each tag for each vTR (**Figure 1B, Supplemental Figure 2**). For each protein, cells expressing the vTR demonstrated a substantially different expression pattern as compared to control cells that only expressed GFP or a puromycin resistance cassette (**Figure 1B** and **Supplemental Figure 2**). The final location of the GFP tag (**Supplemental Figure 3**) was selected based upon similarity of gene expression effects to the untagged vTR using both heatmap-based clustering (**Supplemental Figure 2**) and PCA analysis (**Supplemental Figure 4**). AlphaFold3 (55) predictions affirmed that the selected GFP tag location would likely not disrupt protein folding (**Supplemental Figure 3**).

Once the nine EBV vTRs were selected for further study, we generated isogenic stable cell lines expressing each vTR, with a Flp recombinase enabling integration into a specific genomic location (*FRT*) with the preferred N or C terminal tag, resulting in consistent vTR expression across the cell lines (**Figure 1A, Supplemental Figure 5**, and see Methods). These isogenic stable cell lines were then used to study the impact of EBV vTRs on gene expression (RNA-seq), interactions with the human genome (ChIP-seq), and chromatin accessibility (ATAC-seq) (**Figure 1A**).

### EBV vTRs induce extensive non-redundant changes to human gene expression

We first performed biological triplicate RNA-seq of each cell line with stable EBV vTR expression (see Methods). When assessed together, the resulting data demonstrate consistent expression levels and strong quality control metrics (**Figure 1 C** and **D**). We next performed differential analysis of each vTR against the control cell line expressing GFP. Genes with a fold-change of at least 20% and an adjusted p-value less than 0.05 were called differentially expressed. A 20%-fold-change threshold was chosen based on the identification of an inflection point in the number of differential genes across various thresholds that was consistent across all vTRs (**Supplemental Figure 6**). Of the 61,544 human coding and non-coding genes that we examined, 15,039 (24%) were impacted by at least one EBV vTR. Expression of each of the nine selected EBV vTRs resulted in robust changes to gene expression, ranging from 2,069 to 7,234 differentially expressed genes (DEGs) (**Figure 1C**). Overall, EBV vTRs regulate human gene expression in a balanced fashion: expression of all vTRs results in upregulation and/or downregulation between 45% and 55% of differentially expressed host genes (**Figure 1C**).

Unexpectedly, BMRF1 had a greater impact on host gene expression than any of the eight known EBV vTRs (**Figure 1C**). To our knowledge, only a single study of this protein’s impact on human gene expression has previously been performed (56).

EBV vTRs broadly regulate different host genes (**Figure 1D**), and the biological processes that are enriched for EBV vTR upregulated genes are largely different from those enriched for downregulated genes (**Figure 1E**, **Supplemental Figure 7**). Biological processes enriched among genes that are significantly upregulated by EBV vTR expression were largely shared between the vTRs, and include wound healing, inflammatory response, cytoskeleton organization, and cell adhesion (**Figure 1E** and **Supplemental Figure 7**). By contrast, expression of numerous EBV vTRs led to the downregulation of genes enriched for mitochondrion organization, DNA repair, and chromosome organization (**Figure 1E** and **Supplemental Figure 7**). We note that most mitochondrial proteins are encoded by the host nuclear genome. Collectively, these results indicate that EBV vTRs have widespread effects on a wide range of human genes and pathways.

### EBV vTRs occupy thousands of shared and distinct regions across the human genome

We next sought to systematically catalog the regions of the human genome occupied by each of the nine vTRs that substantially alter human gene expression levels. To this end, we performed anti-GFP ChIP-seq in the nine cell lines harboring each vTR (see Methods). The resulting data were of high quality, meeting or exceeding the standards established by the ENCODE consortium (57, 58). Fraction of Reads in Peaks (FRiP) scores all exceeded 0.03, every replicate set had at least 1,500 reproducible peaks, and the replicates clustered together tightly and distinctly from GFP-only controls (**Supplemental Figure 8**). Further, vTR ChIP-seq peaks were enriched for known DNA binding motifs (ZTA, EBNA1) or known interacting human TF motifs (EBNA2 with human RBPJ motifs). ChIP-seq peaks also recapitulate known biology in other experimental systems, with strong EBNA2 ChIP-seq peaks coinciding with previously published EBNA2 ChIP peaks (18) (**Figure 2A**, middle, **Supplemental Figure 9**). For example, the *MAP3K8* gene is strongly induced by EBNA2 in our system (**Figure 2A**, top), has an EBNA2 ChIP-seq peak in the promoter, and has been shown to be transcriptionally regulated by EBNA2 (17, 26). Additional examples involving known EBNA2-based gene regulatory mechanisms (e.g., *BMF*, *CDKN1A*, *CSRNP1*, and *CXXC5* (17)) are provided in **Supplemental Figure 9.** Taken together, these results indicate that this ChIP experimental system is of high quality and is capable of recapitulating previously established molecular mechanisms.

**Figure 2.**
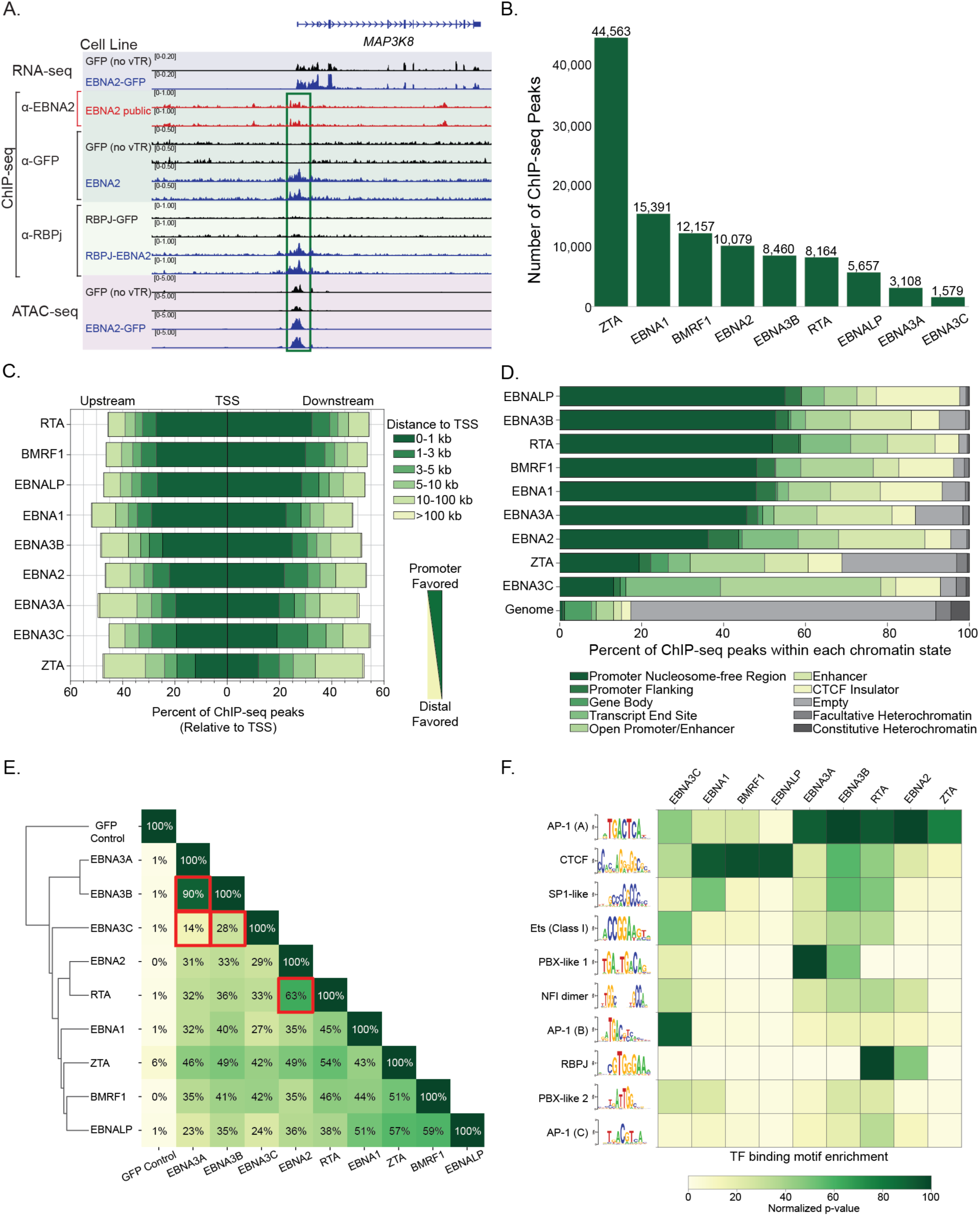
EBV vTRs occupy thousands of shared and distinct regions across the human genome. **A.** Example of EBV vTR-induced human gene regulatory changes. Top rows: EBNA2-induced alteration to *MAP3K8* gene expression (RNA-seq). Vertical green outlined box, top-to-bottom: EBNA2 binding in a public dataset (red) and in our system (ChIP-seq), EBNA2-altered RBPJ binding (ChIP-seq), and EBNA2-altered chromatin accessibility (ATAC-seq). **B.** EBV vTR ChIP-seq peak counts in the human genome. **C.** vTR ChIP-seq peak distribution relative to human transcription start sites (TSS). vTRs are sorted top-to-bottom, in order of “promoter favored” to “distal favored.” **D.** vTR ChIP-seq peak distribution within human chromatin states. Chromatin states were calculated using a variety of genome-scale epigenetic measurements (see Methods). vTRs are sorted top-to-bottom, in order of “promoter associated” to “non-promoter associated” states. The frequency of each state across the human genome is provided at the bottom for reference. **E.** Pairwise overlap of vTR ChIP-seq peaks. Each entry indicates the percent overlap (number of intersecting peaks divided by the number of peaks in the smaller peak set) between the given pair of vTR ChIP-seq datasets. Rows and columns were hierarchically clustered using the Unweighted Pair Group Method with Arithmetic Mean (UPGMA) algorithm. vTR pairs discussed in the text are indicated with red boxes. **F.** Human TF binding site motif enrichment within vTR ChIP-seq peaks. The DNA sequences of vTR ChIP-seq peaks were subjected to motif enrichment analysis (see Methods). The top 10 unique motifs across all nine vTRs are shown. Non-redundant motifs were established through manual curation. Normalized p-values indicate the percentage of the maximum p-value obtained from motif enrichment analysis for each vTR. Motifs are sorted top-to-bottom from strongest to weakest cumulative enrichment.

Encouraged by the high quality of our ChIP-seq data, we next took advantage of our systematic approach to compare various biological attributes of the EBV vTRs. In total, EBV vTRs occupy 64,290 distinct regions of the human genome, comprising 62,340,273 total bases (2% of the entire human genome), consistent with their enormous cumulative effect on human gene expression levels (**Figure 1C**). Overall, ZTA had the largest number of binding events, with over 44,000 reproducible ChIP-seq peaks in the human genome (**Figure 2B**). Surprisingly, the understudied BMRF1 protein had over 12,000 peaks in the human genome (**Figure 2B**), implying a potentially important role in EBV-host interactions. All vTRs bind with high symmetry (upstream vs. downstream) with respect to human transcription start sites (TSS) (**Figure 2C**). Some vTRs prefer binding near promoter regions – for example, >80% of RTA and BMRF1 ChIP-seq peaks fall within 5 kb of a TSS. In contrast, ZTA tends to bind more distal regions, with <60% of its peaks within 5kb of the TSS, consistent with the similarity of its structure and DNA binding motif to human AP-1 TFs, which also tend to occupy enhancers (59).

We next examined the types of functional regions that statistically coincide with vTR binding events in the human genome. To this end, we used previously computed chromatin states obtained by running the ChromHMM method (60) on ChIP-seq data for CTCF and six histone marks, as well as ATAC-seq data (61). Notably, all nine vTRs occupied considerably more regulatory regions of the genome (i.e., “non-empty” and “non-heterochromatin” regions) compared to genomic background (**Figure 2D**), consistent with their roles in gene regulation. Nonetheless, almost 30% of ZTA ChIP-seq peaks coincide with empty or heterochromatin regions, suggesting that it might have similar roles to recently discovered “Dark TFs” (61). Consistent with the TSS analysis, ZTA and EBNA3C both tend to avoid promoters relative to the other vTRs. The binding events of three vTRs (EBNALP, EBNA1, and BMRF1) strongly coincide with CTCF insulators, suggesting possible roles in the alteration of chromatin architecture through CTCF-based mechanisms. Collectively, these data indicate that the binding events of all nine vTRs strongly localize within regulatory regions of the human genome, with specific vTRs likely participating preferentially in different gene regulatory mechanisms.

We next compared the location of vTR binding events relative to one another. Overall, EBV vTRs occupy largely distinct regions of the genome, with a median pairwise ChIP-seq peak overlap of only 39% (**Figure 2E**). The strongest peak overlap (90%) was observed for EBNA3A and EBNA3B, consistent with their known preference to bind to the same regions of the genome (51). In contrast, EBNA3C had minimal overlap with EBNA3A and EBNA3B (14% and 28%, respectively). EBNA2 and RTA peaks displayed strong overlap (63%), which is surprising given that they are expressed in mutually exclusive stages of the EBV life cycle (latent vs. lytic, respectively) (62).

Finally, we examined the enrichment of human TF DNA binding motifs within the vTR-occupied regions of the human genome (see Methods). As expected, ZTA enriched for AP-1-like motifs, and EBNA2 enriched for motifs of its well-established binding partner RBPJ (63) (**Figure 2F**). Surprisingly, RTA peaks displayed even stronger enrichment for RBPJ motifs. To our knowledge, this represents the first evidence that EBV-encoded RTA may engage with RBPJ. The RTA encoded by Kaposi’s Sarcoma-associated Herpesvirus (KSHV) is known to interact with RBPJ (64). However, KSHV-RTA shares only limited sequence homology with its EBV counterpart (25% amino acid identity, 38% similarity). Strikingly, all nine vTRs strongly enriched for either CTCF (EBNA1, BMRF1, EBNALP) or AP-1 (all other vTRs) motifs, but not both. These results are consistent with our chromatin state analyses, which implicate these same vTRs at CTCF insulators and enhancers, respectively (**Figure 2D**). Certain other motifs, such as SP1, Ets, PBX, and NFI, are enriched for multiple vTRs, suggesting that they might be shared EBV vTR targets. In summary, these analyses demonstrate that EBV vTRs occupy over 64,000 distinct regions across the human genome, exhibiting variation in genomic localization, preferences for specific chromatin states, and enrichment for human TF motifs.

### EBV vTR expression robustly changes human chromatin accessibility

We next examined the effect of each vTR on human chromatin accessibility. To this end, we performed ATAC-seq experiments in cells expressing either GFP alone (as a control) or each vTR independently. Quality control analyses indicated that each dataset was of high quality, with FRiP scores of at least 0.15 (median of 0.32), TSS enrichment scores of at least 4 (median of 11.18), and at least 45,000 ATAC-seq peaks (median of 71,112).

We first compared the regions of open chromatin in each of the ten cell lines. All vTRs had at least a moderate impact on human chromatin, with a range of 76% to 93% shared peaks with the GFP control cell line (**Figure 3A**). The lone exception was ZTA, which shared only 63% of its peaks with the control cell line, suggesting that ZTA has potent pioneering-like activity. To further examine these phenomena, we next identified statistically robust chromatin opening and closing events by comparing the ATAC-seq signal strength of each vTR cell line with the GFP control cell line (see Methods). Differential ATAC-seq peaks for each respective vTR were partitioned into peaks that are likely to be directly altered by that vTR and those likely to be impacted by

**Figure 3.**
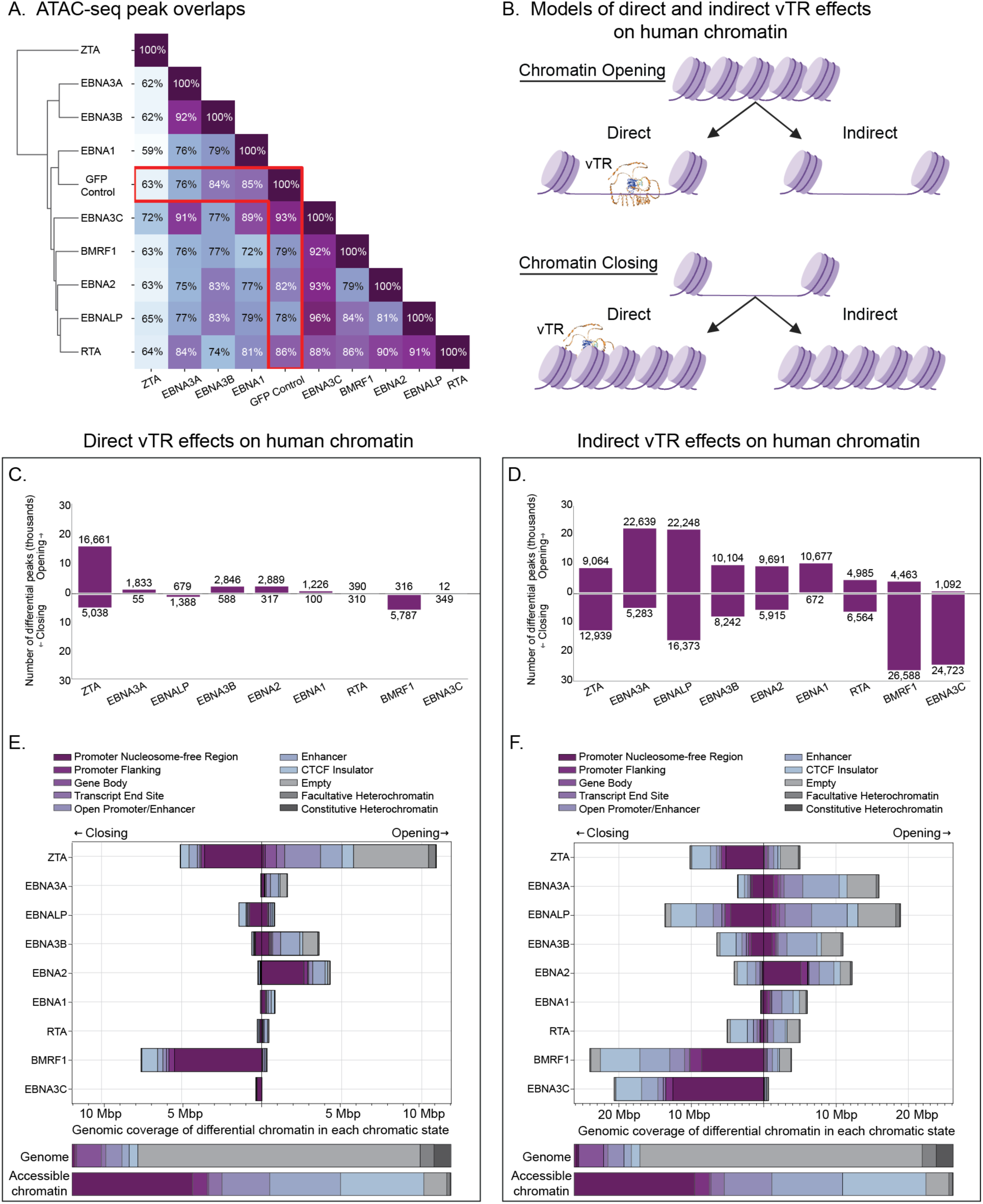
EBV vTR expression robustly changes human chromatin accessibility. **A.** Pairwise overlap of ATAC-seq peaks in vTR expressing cell lines. Each entry indicates the percent overlap (number of intersecting peaks divided by the number of peaks in the smaller peak set) between the given pair of vTR ATAC-seq datasets. Rows and columns were hierarchically clustered using the Unweighted Pair Group Method with Arithmetic mean (UPGMA) algorithm. GFP control, which indicates peak overlap with cells lacking a vTR, is boxed in red. **B.** Models of direct and indirect vTR effects on human chromatin. We define genomic regions with direct opening or closing of chromatin as vTR-induced differential ATAC-seq peaks that directly overlap with a ChIP-seq peak for that vTR. All other differential ATAC-seq peaks are defined as indirect. **C and D.** Quantification of vTR-altered ATAC-seq peaks (C, direct; D, indirect). For each vTR, the total number of differential ATAC-seq peaks are indicated (up bars, opening; down bars, closing). **E and F.** vTR-altered ATAC-seq peak distribution within human chromatin states (E, direct; F, indirect). Chromatin states were calculated using a variety of genome-scale epigenetic measurements (see Methods). The frequency of each state across the human genome and within accessible chromatin is provided at the bottom for reference.

We next sought to characterize the potential functional impact of vTR-induced chromatin accessibility changes. To this end, we compared the genomic locations of vTR-induced chromatin opening and closing events to ChromHMM-calculated chromatin states, again distinguishing between direct and indirect effects. The majority of direct EBV vTR-induced chromatin accessibility changes are located within or near human gene promoters (**Figure 3E**). Specifically, EBNA2 tends to open promoters and BMRF1 and ZTA tend to close promoters (**Figure 3E**). The majority of ZTA-induced direct chromatin opening events do not involve promoters but instead consist largely of empty and heterochromatin regions of the genome, consistent with the preferential binding of ZTA to these locations (**Figure 2D**). Examination of indirect vTR effects further underscores many of these observations, including the predominant effects of EBNA2 opening promoters, ZTA closing promoters, and BMRF1 closing promoters (**Figure 3F**). Although EBNA3C has only limited direct effects on chromatin accessibility (12 and 349 opening and closing events, respectively), it has a substantial number of indirect closing effects (24,723). Collectively, these results reveal that EBV vTRs have substantial and diverse effects on various regions of the human genome, with differential effects on promoters, distal regions, and empty regions of “genomic dark matter” (61).

Finally, to identify the human TFs that are most strongly impacted by these chromatin changes, we performed TF binding motif enrichment analysis in the direct vTR-induced chromatin accessibility regions. This analysis revealed that AP-1 is the first or second most strongly enriched opened motif for all vTRs, and CTCF is the first or second closed motif for almost every vTR (**Figure 4**). EBNA1 represents an interesting exception by instead strongly opening CTCF motifs and strongly closing Ets motifs. EBNA2, ZTA, and EBNA1 uniquely close AP-1 sites among vTRs. RBPJ motifs are opened by both EBNA2 and RTA, consistent with their strong enrichment within the ChIP-seq peaks of these proteins (**Figure 2F**). Intriguingly, two different classes of homeoboxes are specifically enriched within the opened regions of EBNA3A/3B (PBX-like) and EBNA3C (NKX-like) (**Figure 4**). Collectively, these results indicate that EBV vTRs induce extensive alterations to human chromatin accessibility that primarily impact promoters through a variety of mechanisms (opening vs closing, various impacted classes of human TFs).

**Figure 4.**
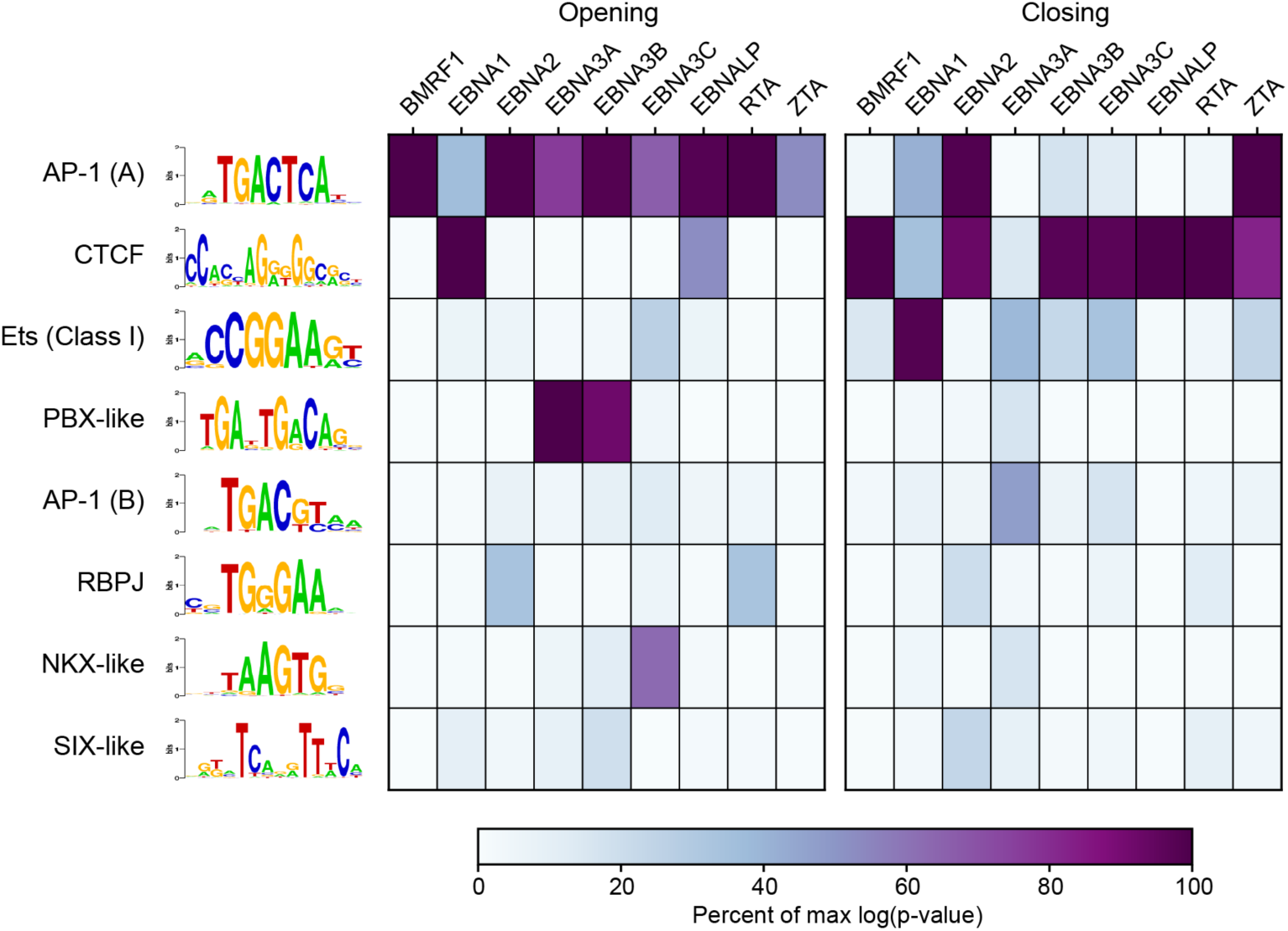
Human TF motif enrichment in regions of the human genome with chromatin accessibility directly altered by EBV vTRs. ATAC-seq experiments were performed in human cells following the expression of one of nine vTRs or GFP as a control. These data were used to identify vTR-induced alterations to chromatin accessibility across the human genome (see Methods). Human TF motif enrichment analysis was performed in these opening or closing chromatin regions for each vTR. Normalized p-values indicate the percentage of the maximum p-value obtained from motif enrichment analysis for each vTR, using a curated non-redundant set of motifs that are strongly enriched for at least one vTR.

### An integrated assessment of EBV vTR interactions with the human genome

We next sought to examine the relationships across data types for each vTR. We first asked if vTR induced alterations to human gene expression (RNA-seq) coincided with vTR binding to the human genome (ChIP-seq). Indeed, for each vTR, we observed strong enrichment of ChIP-seq peaks within a 10kb region of genes induced by that vTR. Binding of each vTR had significant enrichment for both upregulated and downregulated genes (10^-190^ < P < 10^-5^, up to 5-fold enriched) (**Supplemental Figure 10**, panel A). Likewise, binding of each vTR had significant enrichment for vTR-induced opening and closing chromatin regions (10^-215^ < P < 0.02, up to 16-fold enriched) (**Supplemental Figure 10**, panel B).

Data integration also reveals multiple compelling examples of EBV vTR-altered gene regulatory mechanisms. As an example, we highlight six regions (**Supplemental Figure 9**) in which EBNA2 binds, increases chromatin accessibility, and induces gene expression of genes that are involved in the interferon response to viruses or are known to be regulated by EBNA2 (65–70). At each locus we also observe RBPJ binding in our system and EBNA2 binding in B cells with whole virus infection (18, 71). Collectively, these results indicate a strong concordance between the data types obtained in this study.

### EBNA2 and RTA independently stabilize RBPJ DNA binding to the human genome

Many vTRs exert their regulatory functions through interactions with host factors (23). For example, EBNA2 is a well-established Notch mimic that interacts with and stabilizes RBPJ binding to the host and viral genomes (63). In our integrative analyses, we uncovered several lines of evidence suggesting that a second EBV-encoded protein, RTA, may also interact with RBPJ. First, EBNA2 and RTA ChIP-seq peaks have substantial overlap (63%, ranked second behind the EBNA3A/EBNA3B pair) (**Figure 2E**). Second, RTA ChIP-seq peaks are enriched for RBPJ motifs even more strongly than EBNA2 ChIP-seq peaks (**Figure 2F**). Third, differentially accessible human genomic regions in the presence of RTA are strongly enriched for RBPJ motifs, similar to what we observe for EBNA2 (**Figure 4**). Fourth, the top enriched *de novo* motif in RTA ChIP-seq peaks is a near-perfect match to the known RBPJ motif (**Figure 5A**). To further examine the possibility that RTA and RBPJ may interact, we performed ChIP-seq for RBPJ in human cells expressing EBNA2, RTA, or GFP only (as a control). The resulting data met our high quality control standards, with FRiP scores exceeding 0.09, peak counts above 20,000, very strong enrichment for RBPJ motifs, and replicates clustering together tightly and distinctly from GFP-only controls (**Supplemental Figure 11**).

**Figure 5.**
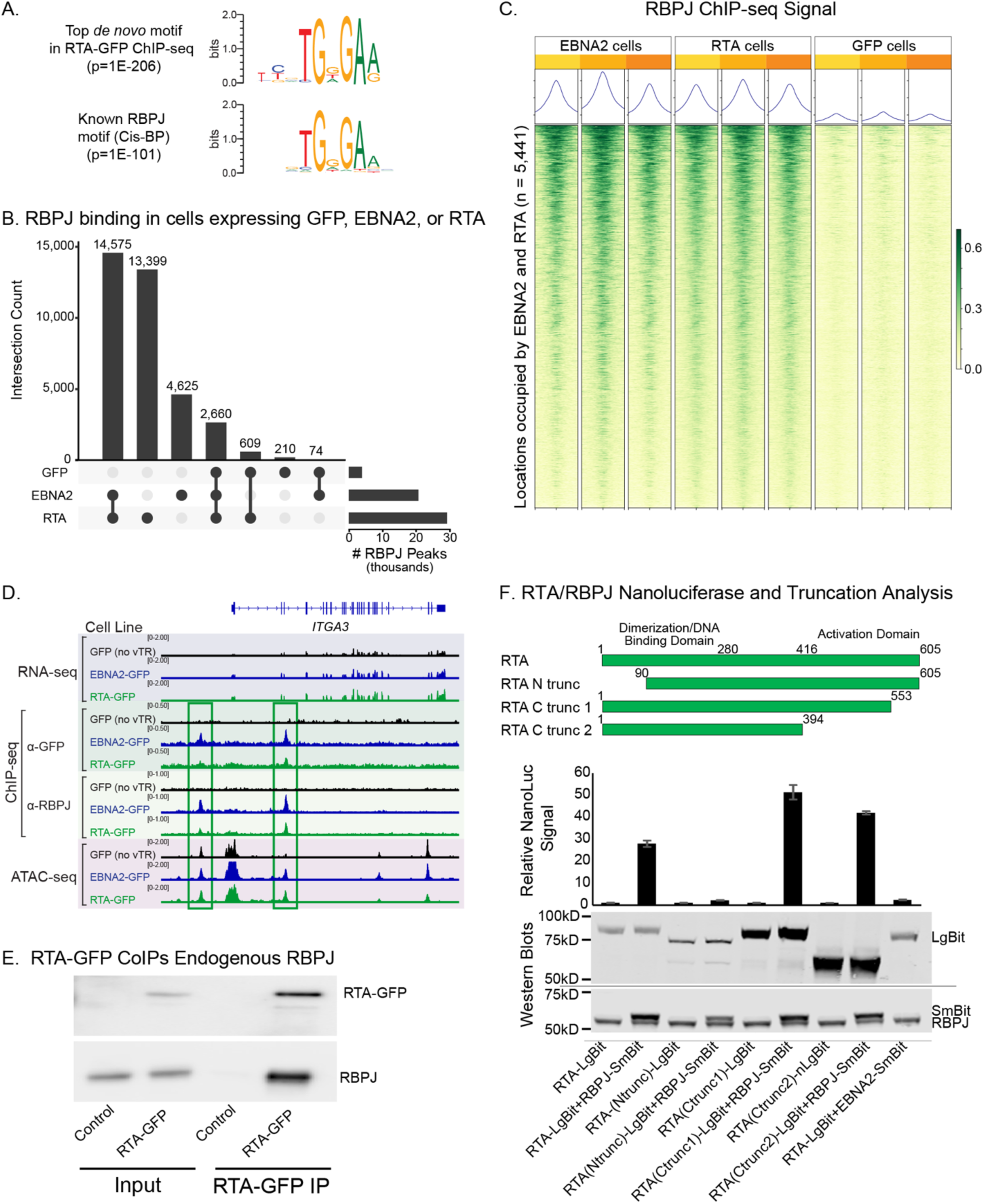
EBNA2 and RTA independently stabilize RBPJ DNA binding to the human genome. **A.** RTA-GFP ChIP enriches the RBPJ motif. Top: #1 *de novo* enriched motif obtained from RTA ChIP-seq peaks. Bottom: the known RBPJ DNA binding motif. **B.** RBPJ ChIP-seq peak overlap counts in cells expressing GFP (control), EBNA2, or RTA. **C.** Normalized RBPJ ChIP-seq signal in EBNA2, RTA, or GFP (control) cells at sites ChIPed by both EBNA2-GFP and RTA-GFP. Values indicate counts per million. Window size is –1Kb to 1Kb for each column. **D**. Genome browser screenshot of the *ITGA3* gene, which is upregulated in both EBNA2 and RTA expressing cells. Track sets show (top-to-bottom): RNA-seq, anti-GFP ChIP-seq, anti-RBPJ ChIP-seq, and ATAC-seq signal. For each data type, data are presented in cells expressing (top-to-bottom): GFP, EBNA2, or RTA. Green boxes indicate genomic regions discussed in the text. **E**. Co-immunoprecipitation of RTA-GFP with endogenous RBPJ. Lane order (left to right): input from control cells, input from RTA-GFP cells, GFP immunoprecipitation from control cells, GFP immunoprecipitation from RTA-GFP cells that were probed for RTA-GFP (top blot) or RBPJ (bottom blot). **F**. Split NanoLuc luciferase assays examining the interaction between RTA and RBPJ. Top: Schematic of RTA protein and RTA protein truncations. Middle: Luciferase signal obtained from cells expressing the indicated proteins. Bottom: Western blots on the same cells. Note the similar expression levels of the RTA-LgBit and the RTA truncations, equivalent expression levels of RBPJ-SmBit across cells, and similar endogenous RBPJ expression levels.

We next compared the binding profiles of RBPJ in cells expressing GFP, EBNA2, or RTA. Notably, RBPJ binding is greatly increased in the EBNA2- and RTA-expressing cells (22,978 and 32,130 peaks, respectively) compared to the GFP-only condition (3,744 peaks) (**Figure 5B**). RBPJ’s enhanced binding with EBNA2 and RTA is likely due to stabilization, as RBPJ binds transiently when it lacks a cofactor such as Notch (72) or EBNA2 (73). Further, the largest class of RBPJ ChIP-seq peaks represents places where RBPJ binds the human genome both in the presence of EBNA2 and in the presence of RTA, but not in the presence of GFP (**Figure 5B**). Indeed, the presence of either EBNA2 or RTA leads to consistently stronger RBPJ ChIP-seq signal within regions at which both EBNA2 and RTA independently localize (**Figure 5C**). An example genomic region is shown in **Figure 5D** for the *ITGA3* gene, which is a known RBPJ target (74). *ITGA3* gene expression levels are significantly increased relative to GFP controls with expression of EBNA2 or RTA (**Figure 5D**, top). Two regions proximal to the *ITGA3* TSS contain binding events for both EBNA2 and RTA that coincide with RBPJ binding events that only occur in the presence of EBNA2 or RTA (and not in the presence of GFP only) (**Figure 5D**, middle). The intronic region also displays both EBNA2-and RTA-dependent chromatin accessibility, while the promoter region does not (**Figure 5D**, bottom). Additional examples of EBNA2- and RTA-dependent RBPJ binding are provided in **Supplemental Figure 12**. Collectively, these results reveal that both EBNA2 and RTA likely stabilize RBPJ binding at thousands of regions of the human genome.

To investigate whether RTA and RBPJ proteins physically interact, we conducted a series of biochemical assays. To this end, we first confirmed that RTA directly interacts with endogenous RBPJ through co-immunoprecipitation assays (**Figure 5E**). Notably, the amount of RBPJ does not change in cells stably expressing RTA compared to control cells (**Figure 5E**, bottom left), indicating that the massive increase in RBPJ binding events in RTA-expressing cells (**Figures 5B** and **5C**) is not simply due to increased amounts of available RBPJ protein. To further characterize this interaction, we employed split NanoLuc luciferase assays, wherein the two NanoLuc luciferase halves (SmBit and LgBit) were fused to (1) RBPJ and (2) one of several RTA forms, including full length RTA and various truncated forms (**Figure 5F**). Co-expression of RTA-LgBit and RBPJ-SmBit resulted in a significant increase in luminescence compared to RTA-LgBit alone, confirming an interaction between the two proteins. (**Figure 5F**). Analysis of the truncated forms of RTA revealed that a small N-terminal truncation (aa 1-89) in the region containing the RTA DNA binding domain significantly reduced luciferase signal. In contrast, deletions of C-terminal regions (amino acids 554– 605 and 395–605) did not impair the interaction (i.e., the signal was retained). Collectively, these results confirm that RTA colocalizes with RBPJ on the human genome and reveal that the RTA DNA binding domain is necessary for this colocalization.

### EBV BMRF1 regulates mitochondrion organization genes with enriched binding at Rheumatoid Arthritis genetic risk loci

Through a comprehensive analysis of BMRF1’s impact on human gene regulation, this study offers a valuable first glimpse into the function of a largely uncharacterized EBV vTR. Of all EBV vTRs examined, BMRF1 had the largest impact on gene expression (**Figure 1C**), with 1,396 genes that are uniquely regulated by this protein. Notably, the majority of these genes were downregulated in the presence of BMRF1. Integrated multi-omic data support a model in which BMRF1 generally binds to promoters, closes chromatin, and reduces gene expression levels (**Figure 2**, **Figure 3**, **Supplemental Figure 10**).

As little is known about this protein, we focused our efforts on deeper characterization of the genes and pathways impacted by BMRF1. Our data implicate an important role for BMRF1 in the reduction of mitochondrial organization gene expression levels (**Figure 1E**). In total, 209 of 589 genes involved in this process are regulated by BMRF1 (**Figure 6A**). As an example, *NDUFS7* expression levels are reduced with BMRF1 promoter binding, with a concomitant decrease in chromatin accessibility (**Figure 6B**). NDUFS7 is a component of the “mitochondrial complex I”, with mutations in the protein implicated in Leigh Syndrome, a rare neurodegenerative disorder (75, 76). There are many additional examples of mitochondrial organization genes with similar BMRF1-based mechanisms (**Supplemental Figure 13**).

**Figure 6.**
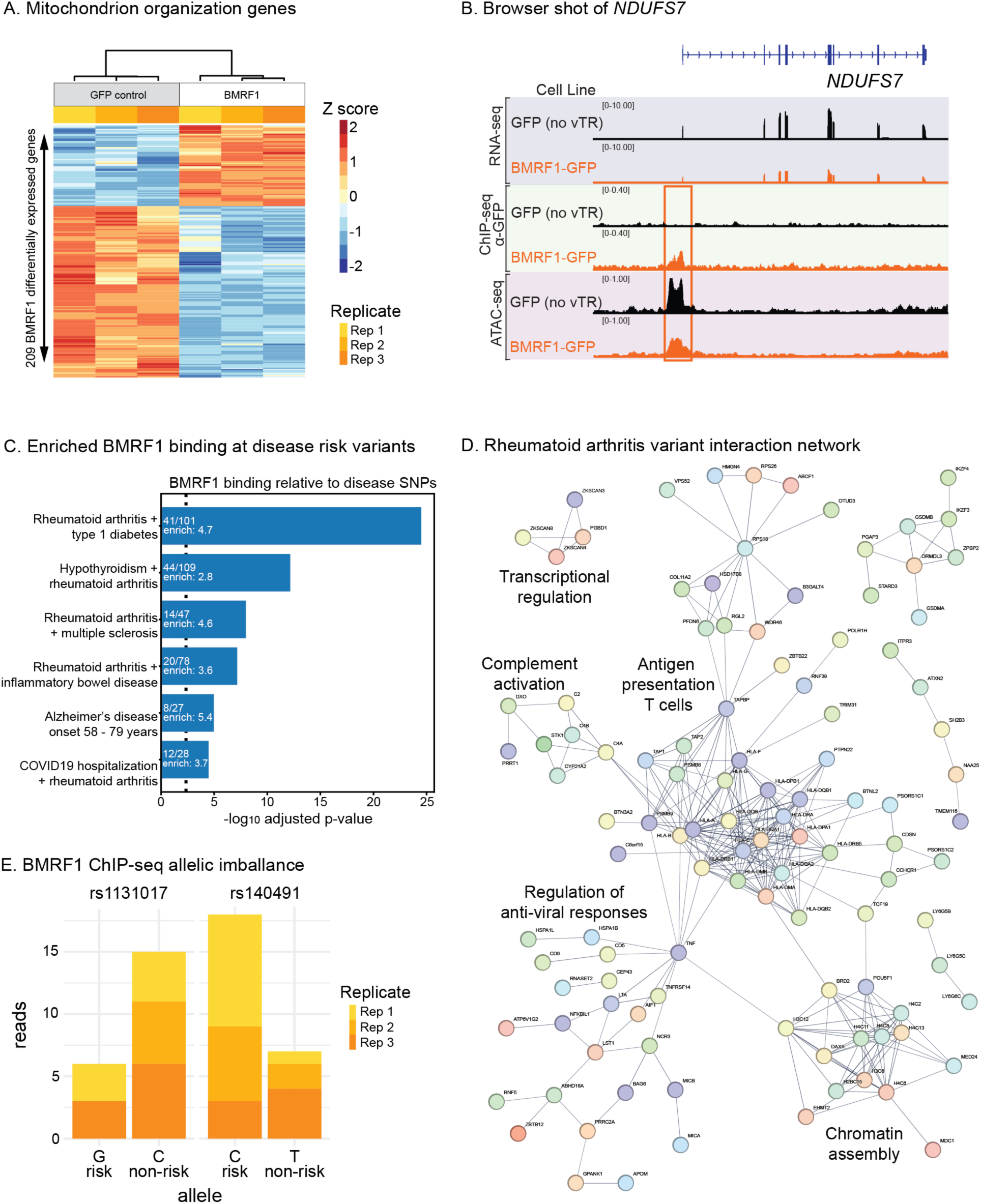
BMRF1 regulates mitochondrion organization genes, with enriched binding at rheumatoid arthritis genetic risk loci. **A.** Quantification of BMRF1 induced differential genes in the mitochondrion organization process in cells stably expressing BMRF1 or a GFP control. Values indicate normalized gene expression levels that have been Z-transformed across rows (replicates). Columns and rows were clustered using Ward’s method. **B.** Genome browser screenshot of the *NDUFS7* gene, which is downregulated and shows reduced promoter chromatin accessibility with BMRF1 expression. Track sets show (top-to-bottom): RNA-seq, anti-GFP ChIP-seq, and ATAC-seq signal. For each data type, data are presented in cells expressing (top-to-bottom) GFP or BMRF1-GFP. Orange box indicates the promoter region discussed in the text. **C.** Enrichment of BMRF1-GFP ChIP-seq peak sets at disease risk loci. All assessed datasets with significant enrichment (adjusted p-values less than p = 0.01) are shown. The –log10 adjusted p-value is graphed on the x-axis and the enrichment is provided within each bar. A dotted line indicates the threshold for statistical enrichment. **D.** STRING representation of genes linked to BMRF1-GFP ChIP-seq peaks at rheumatoid arthritis (RA) risk loci, as identified using the eQTLs (98). Connected gene clusters were labeled based on pathway enrichment. **E.** Allelic BMRF1-GFP ChIP-seq at two RA risk variants as assessed using the MARIO tool (see Methods). The number of reads corresponding to the reference and non-reference allele are quantified for each replicate.

Our work and others have established an important role for the binding of EBV vTRs at genetic risk loci that mediate chronic human diseases (16, 18, 19, 23, 77, 78). Building on this foundation, we conducted an unbiased, systematic analysis using the GWAS Catalog (see Methods, (79)) and found that BMRF1 binding events are highly enriched at genetic risk variants for a range of diseases, including rheumatoid arthritis, type 1 diabetes, hypothyroidism, multiple sclerosis, inflammatory bowel disease, Alzheimer’s disease, and SARS-CoV-2 hospitalization (**Figure 6C**). Many genetic risk loci are shared between diseases, and some GWAS focus on these pleotropic risk loci. Impressively, BMRF1 was implicated at rheumatoid arthritis loci in five out of the six GWAS with significant BMRF1 enrichment (**Figure 6C**). When these BMRF1-bound variants are connected to genes through expression quantitative trait loci analysis (see Methods), a network emerges of interacting genes that impact processes implicated in the pathoetiology of rheumatoid arthritis, including complement activation (80), antigen presentation to T cells (81), chromatin assembly (82), and anti-viral responses (83) (**Figure 6D**). The role of BMRF1 in mitochondrial biology might have direct implications on RA, as mitochondria are an important source of intracellular reactive oxygen species, and facilitate the activation of the inflammasome (84), which produces cytokines linked to disease symptoms in rheumatoid arthritis (85, 86).

Allele-dependent biology plays an important role in diseases with genetic components. When a genetic variant is heterozygous, both alleles are present in cells, allowing allele-dependent measurements. We used the MARIO computational tool (16) to identify allelic BMRF1 binding events to RA variants in our ChIP-seq data (see Methods). These analyses revealed two compelling allele-dependent BMRF1 binding examples at risk variants within a single rheumatoid arthritis risk locus (**Figure 6E**). Notably, one of these variants (rs140491) has previously been shown to result in allele-dependent reporter activity and allele-dependent CTCF binding in EBV infected B cells (87), suggesting a possible disease mechanism involving BMRF1 and CTCF at this locus. Collectively, these results implicate BMRF1 in rheumatoid arthritis-based genetic mechanisms.

### EBV vTRs consistently impact a coordinated network of host genes

Finally, we sought to obtain a systems level understanding of the complex EBV vTR human genome interaction landscape. To this end, we created a network linking each vTR to its likely human gene targets using gene expression and vTR binding data (see Methods) (**Figure 7A**). This network reveals that a substantial number of human genes (2,438) are regulated by multiple EBV vTRs. To further investigate this phenomenon, we examined the expression patterns of all human genes regulated by five or more vTRs (**Figure 7B**). Strikingly, the vast majority of these genes can be classified as either consistently upregulated (53%) or downregulated (31%) across vTRs, but not a mixture of the two (only 16%). The most heavily targeted genes (seven or more vTRs) include *SOCS3* (mixed regulation), *BCL11B (*down regulation), and *SMAD7* (up regulation). Dysregulation of SOCS3 function has been implicated in numerous chronic disease types including allergy, autoimmunity, and cancer (88). BCL11B controls cellular proliferation and promotes apoptosis in the context of viral infection (89). SMAD7 promotes the differentiation of numerous immune cells and has anti-inflammatory effects (90, 91); additionally, SMAD7 has roles in epithelial cell differentiation, which is directly involved in the EBV lytic cycle (92). Consistent with its upregulation by most EBV vTRs, SMAD7 expression is increased in viral infection (93). The interaction network of genes regulated and bound by five or more EBV vTRs also illustrates the connectivity of consistently regulated genes (**Figure 7C**). Intriguingly, antiviral effectors STAT1 and STAT3 are consistently upregulated. Collectively, our data reveal a vast human gene regulatory network coordinately regulated by a complex system of EBV vTR interactions with the human genome.

**Figure 7.**
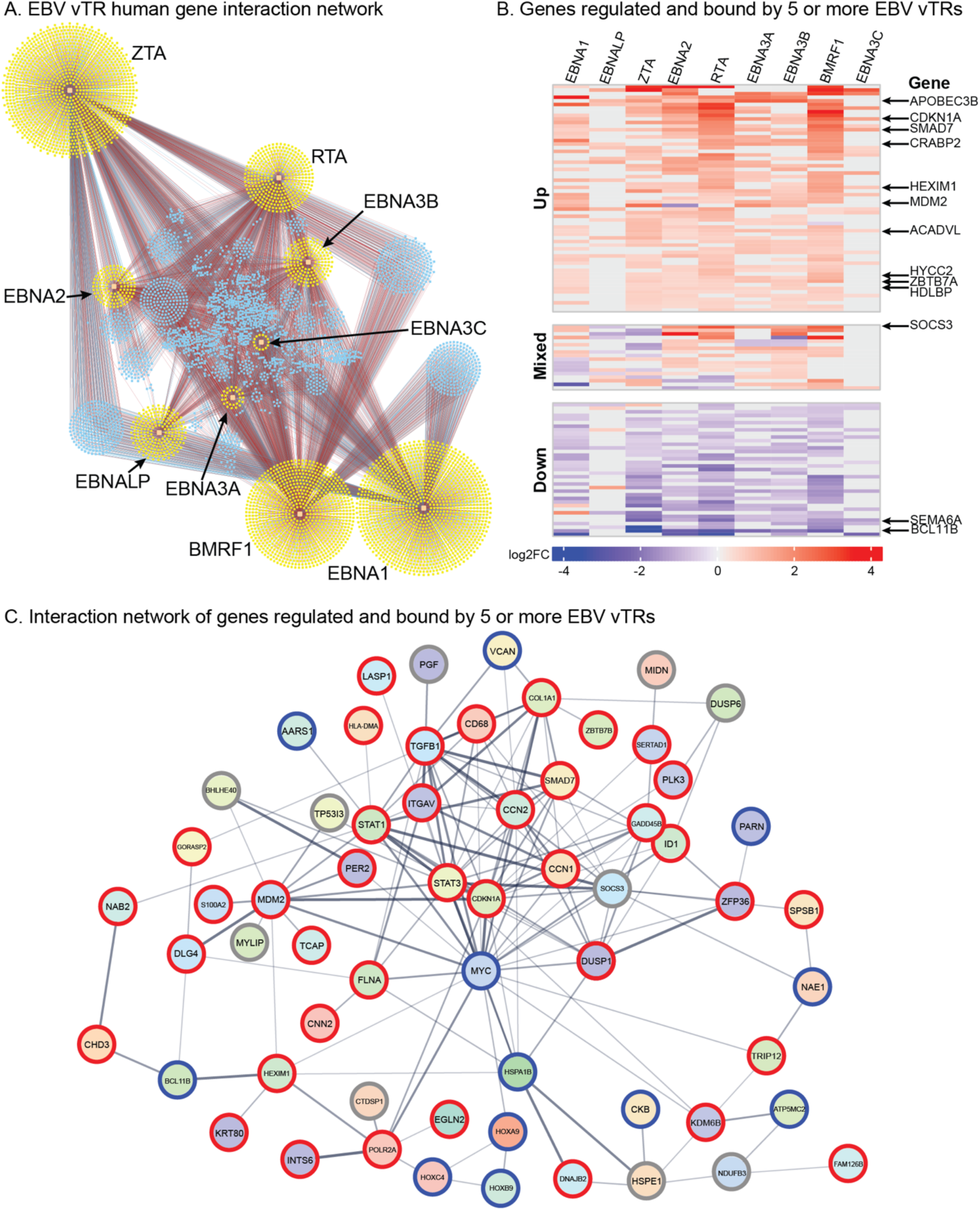
EBV vTRs consistently impact a coordinated network of host genes. **A.** Network representation of genes regulated by vTRs. Nodes indicate vTRs (large squares) or human genes (small squares) with vTR dependent gene expression and a proximal ChIP-seq peak (within 5 kb of TSS) for that vTR. Edges connect vTRs to regulated genes. Edges are colored based on the direction of differential expression (red = up, blue = down). Yellow genes are regulated by a single vTR. Blue genes are regulated by multiple vTRs. **B.** Heatmap of genes (n=118) regulated by five or more vTRs. Colors indicate log2 gene expression fold change. Gray cells indicate that the gene was not differentially expressed and bound by the vTR. Genes were initially k-means clustered (k=3), then hierarchically clustered using Ward’s method within each k-means cluster. Columns were clustered using Ward’s method. Clusters are named according to the direction of differential expression. Genes regulated by at least seven vTRs are indicated. **C.** Network showing interactions between the genes represented in panel B obtained from the STRING database. Colored rings indicate membership in the three clusters displayed in panel B (red=up, blue=down, gray=mixed).

### A new resource for EBV-induced alterations to human gene regulation

The data produced in this study represent a novel and unique resource for examining the human gene regulatory mechanisms impacted by EBV vTRs. In total, our study contains 139 RNA-seq, 30 ATAC-seq, and 38 ChIP-seq datasets. To facilitate access to this massive resource, we have made the data available through three freely-available sources: 1) The Gene Expression Omnibus (GEO) database (GEO accession: XXX); 2) A UCSC Genome Browser (94) session (XXX); 3) A project-specific web page for easy browsing of the available datasets and easy download of the functional genomics data produced in this study, in a variety of formats (XXX). Constructs for the tagged and untagged EBV vTRs presented in this study have been deposited in Addgene. We anticipate this unprecedented resource will be of high value to the virology, bioinformatics, functional genomics, and disease genetics communities.

## Discussion

In this study, we conducted the first systematic genome-scale analysis of EBV vTRs. Our experimental system allowed us to compare results across vTRs with isogenic expression, providing well-controlled conditions that allow us to assess causality. As anticipated, we observed binding at many previously characterized regions of the genome by various vTRs. For example, EBNA2 enriched for the motif of RBPJ, its known DNA binding partner, and bound near its known target *MAP3K8* (17, 26). Similarly, ZTA enriched its known motif and exhibited ChIP-seq peaks near established targets such as *FOSB, PDYN* and *SCIMP* (*49*). Additionally, we observed robust chromatin changes induced by EBNA2 and ZTA expression, contextualizing our findings within the broader landscape of EBV functional genomics (16–18, 42–52).

Our data reveal extensive binding of EBV vTRs across the human genome, with substantial direct and indirect effects on human chromatin accessibility and gene expression. Each vTR regulates a different set of genes. Notably, upregulated genes are enriched in consistent pathways, as are downregulated genes, with minimal overlap between the two. This suggests that the virus has evolved to consistently up- or down-regulate particular biological processes. We also found that RBPJ genomic binding is stabilized by both EBNA2 and RTA, with many shared binding locations despite the distinct expression windows of EBNA2 and RTA in latency and lytic phases of EBV infection, respectively. Our findings thus suggest a “hand-off” mechanism between these proteins facilitated by RBPJ, deepening our understanding of the strategies used by EBV to regulate human genes. A related virus, Kaposi Sarcoma Herpes Virus (KSHV), also expresses an RTA protein that is known to interact with RBPJ (95). Another vTR, BMRF1, emerged from this study as a potent regulator of human genes, occupying promoters, closing chromatin, and lowering gene expression for numerous genes. BMRF1 disease implications are profound, with enriched and allelic binding at genetic risk variants of rheumatoid arthritis.

Upon assessing the genomic sequences occupied by vTRs, we generated models of human transcription factors with which EBV vTRs might bind or compete. For example, places where vTRs occupied and opened chromatin were strongly enriched for AP-1 motifs, while loci bound and closed were enriched for CTCF. We further found that all nine vTRs’ genomic binding events are enriched for either CTCF motifs (EBNA1, BMRF1, EBNA-LP) or AP-1 motifs (all others), but not both. When we assessed the context of EBV vTR genomic occupation, each of the vTRs, and especially ZTA was found to occupy the so called “dark matter” regions of the genome. These enigmatic regions have recently been shown to be occupied by the equally enigmatic human C2H2 zinc finger transcription factors (61).

A previous group identified important viral reprograming of the transcription in the lytic cycle that was driven by a viral pre-initiation complex of many of the less characterized vTRs that were also assessed in this study (96). The complex binds a TATT[T/A]AA, TATA box-like sequence and activates transcriptional initiation across tens of thousands of host genes. We only assess one EBV vTR at a time, which is a limitation of our study. While we did not pursue all of the less characterized EBV vTRs we initially tested, we did measure some transcriptional changes from each of them. It is likely that many of the previously described impacts of these proteins are only realized when they come together in a complex.

Our findings have significant implications for understanding how EBV and other viruses impact the human gene regulatory landscape. By linking changes to specific vTRs, we provide a detailed map of EBV regulatory protein interactions with the human genome. Our network analysis elucidates that individual EBV vTRs regulate many human genes, painting a portrait of a complex regulatory landscape. The consistency in gene regulation and directionality across vTRs for a small core set of human genes is striking, highlighting the vast coordinated impact of EBV on the human genome.

In conclusion, this study represents a major advancement in the field of viral genomics, providing holistic insights into the regulatory mechanisms of EBV vTRs. Similar approaches applied to other less well-characterized viruses will also likely yield robust insights and discoveries. Our study offers a new resource for examining EBV-induced alterations to human gene regulation, with experimental reagents and data available and cross referenced on multiple platforms. These data will enable future research in virology, bioinformatics, functional genomics, and disease genetics, paving the way for new discoveries and therapeutic strategies.

## Acknowledgements

This research was funded by National Institutes of Health (NIH) R01 HG010730, U01 AI130830, P01 AI15058501, and U24 HG013078 to M.T.W.; R01 AR073228, R01 NS099068, and R01 AI024717 to M.T.W. and L.C.K.; R01 AI148276, U01 HG011172, and U19 AI070235, and P30 AR070549 to L.C.K.; R01 AI164709 and DE033907 to B.E.G.; T32 AR069512 to P.J.D. We would like to acknowledge the assistance of Cincinnati Children’s Hospital Medical Center Genomics Sequencing Facility (RRID SCR_022630) for high-throughput sequencing services, Ally Yang for experimental advice, Kevin Ernst for computational support, and the members of the Kottyan and Weirauch labs for constructive feedback and discussion.

## Author contributions

Conceptualization: LCK, MTW; Formal analysis: PJD, LEE, SP, KCMFV, XC; Funding acquisition: LCK, MTW; Investigation: MRH, AAD, SHJ, OD, CY, KAD, CF, HKH, OEG; Methodology: MRH, LCK, MTW; Resources: RR, BEG, BZ, TRH; Software: AV; Supervision: MRH, LCK, MTW; Visualization: PJD, MRH, LEE, SP, LPL; Writing (original draft): PJD, MRH, LEE, CRF, LCK, MTW; Writing (review & editing): all authors.

## Supplemental Figures

**Supplemental Figure 1.** Transcriptional assessment of less characterized EBV vTRs using transient expression of untagged proteins. **A.** All human genes with statistically reproducible differential expression induced by any vTR are shown. Values indicate normalized gene expression levels that have been Z-transformed across rows (conditions). Columns and rows are clustered using Ward’s method. **B.** Number of differentially expressed human genes in cells transiently expressing each untagged vTR. **C.** Principal component analysis of genes with statistically reproducible gene expression by any vTR.

**Supplemental Figure 2.** Differential gene expression from transient tests for EBV vTRs with and without GFP tags. For EBNALP, we only have the N-terminus construct because we were unable to successfully generate a construct with the C-terminus tag. For each vTR, all human genes with statistically reproducible differential expression induced by the untagged vTR are shown. Values indicate normalized gene expression levels that have been Z-transformed across rows (conditions). Columns and rows are clustered using Ward’s method.

**Supplemental Figure 3.** AlphaFold3 (97) renderings of vTRs with and without a GFP tag. The coding sequences of the untagged and GFP fusions (N- and C-terminus) were submitted to the AlphaFold3 server to produce the predicted structures on the left of each panel. The confidence of the structure at each position is indicated by a dark green color in the chart on the right for each panel.

**Supplemental Figure 4.** Principal component analysis of human gene expression from transient tests for EBV vTRs with and without GFP tags. For each vTR, all human genes with statistically reproducible differential expression induced by the untagged vTR are shown.

**Supplemental Figure 5.** Western blot confirmation of expected protein sizes. Confirmation that GFP fusion expression in stable cell lines produces proteins of the expected sizes for the various fusions, along with confirmation of equivalent protein loading through reprobing the blot with an antibody against beta-actin.

**Supplemental Figure 6.** Data driven determination of gene expression fold change thresholds. For each vTR, the number of differentially expressed genes was assessed at the indicated fold change. A clear inflection point emerges at 20% that is consistent across all examined vTRs.

**Supplemental Figure 7.** Complete results of functional annotations enriched among up and down regulated genes across cell lines stably expressing each vTR. Enriched pathways were identified using ToppGene (see Methods). Up to the top five most significant GO Slim Biological Process categories are included for each vTR.

**Supplemental Figure 8.** Overlap analysis of individual vTR ChIP-seq replicates. Each entry indicates the percent overlap (number of intersecting peaks divided by the number of peaks in the smaller peak set) between the given pair of vTR ChIP-seq replicates. Peak sets for each replicate contained peaks with q-value < 0.05. Rows and columns were hierarchically clustered using the Unweighted Pair Group Method with Arithmetic mean (UPGMA) algorithm.

**Supplemental Figure 9.** Genome browser screenshots showing examples of EBNA2 binding in our vTR system and in EBV infected B cells at select genes with EBNA2-dependent expression.

**Supplemental Figure 10.** Integration of vTR binding data (ChIP-seq) with vTR induced gene (RNA-seq) and chromatin (ATAC-seq) changes. **A.** Enrichment of overlap between ChIP-seq and up / down regulated genes (within 10 kilobases of the transcription start site). Enrichment for each vTR was assessed using RELI (see Methods). **B.** Enrichment of overlap between ChIP-seq peaks and opening / closing ATAC-seq peaks using RELI.

**Supplemental Figure 11.** Overlap analysis of individual αRBPJ ChIP-seq replicates. Each entry indicates the percent overlap (number of intersecting peaks divided by the number of peaks in the smaller peak set) between the given pair of αRBPJ ChIP-seq replicates. Peak sets for each replicate contain peaks with q-value < 0.05. Rows and columns were hierarchically clustered using the Unweighted Pair Group Method with Arithmetic Mean (UPGMA) algorithm

**Supplemental Figure 12.** Genome browser screenshots of select genes whose expression is upregulated in both EBNA2 and RTA expressing cells. Each example exhibits overlapping binding events for EBNA2 and RTA, along with enhanced RBPJ binding in both EBNA2 and RTA expressing cells.

**Supplemental Figure 13.** Genome browser screenshots of select genes whose expression is altered by BMRF1, along with closing chromatin and BMRF1 binding at the promoter.

## STAR★Methods

### Key resources table

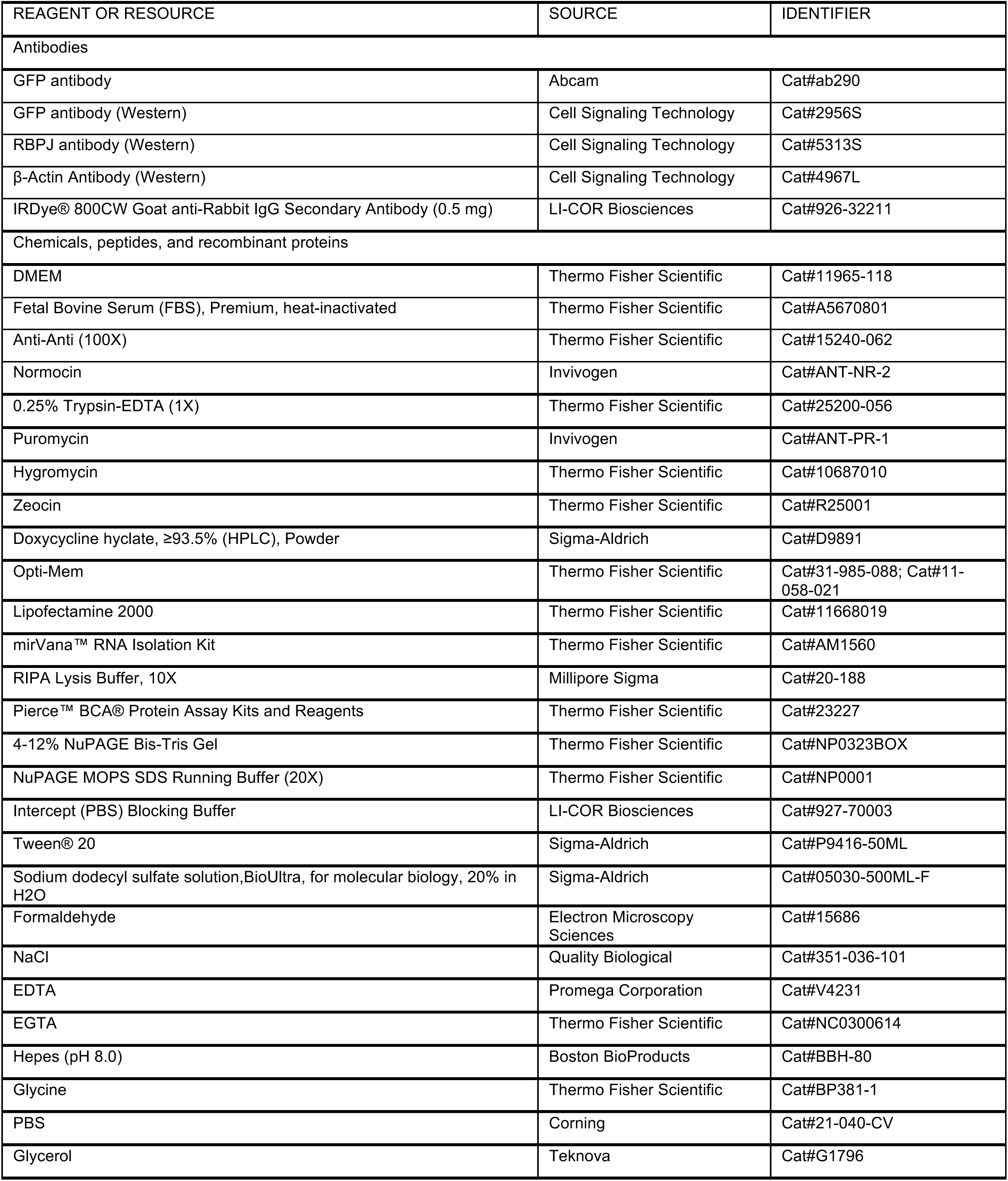
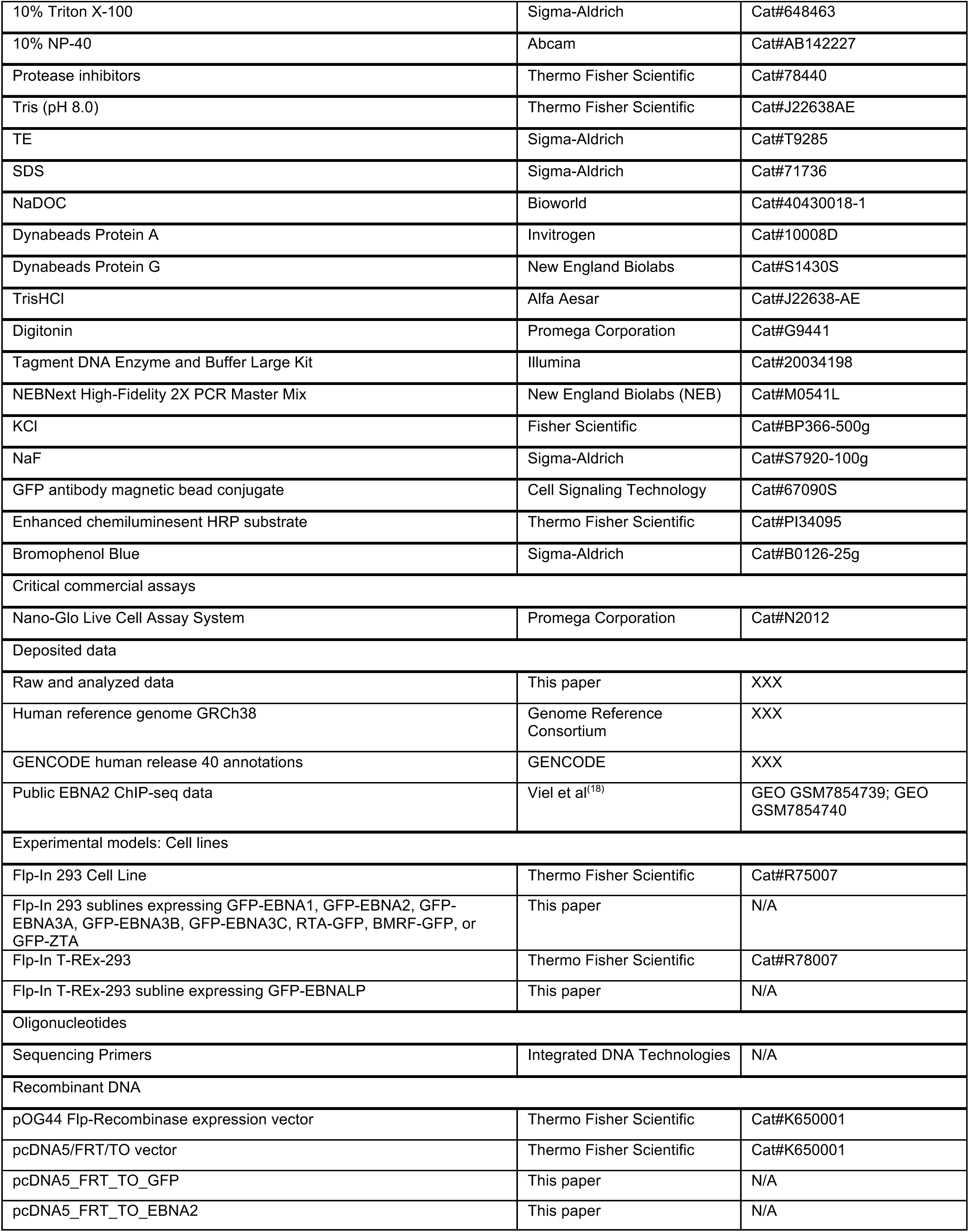
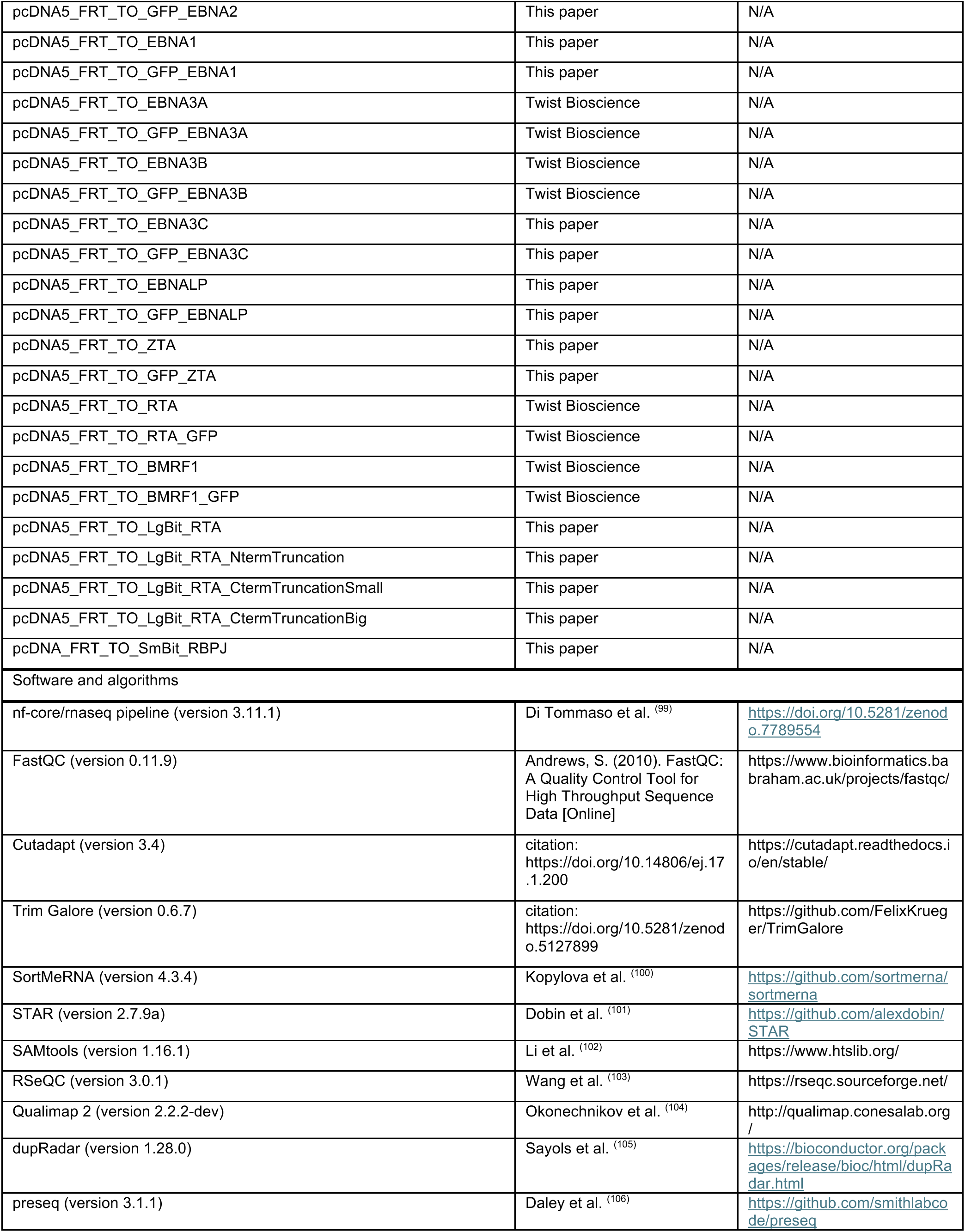
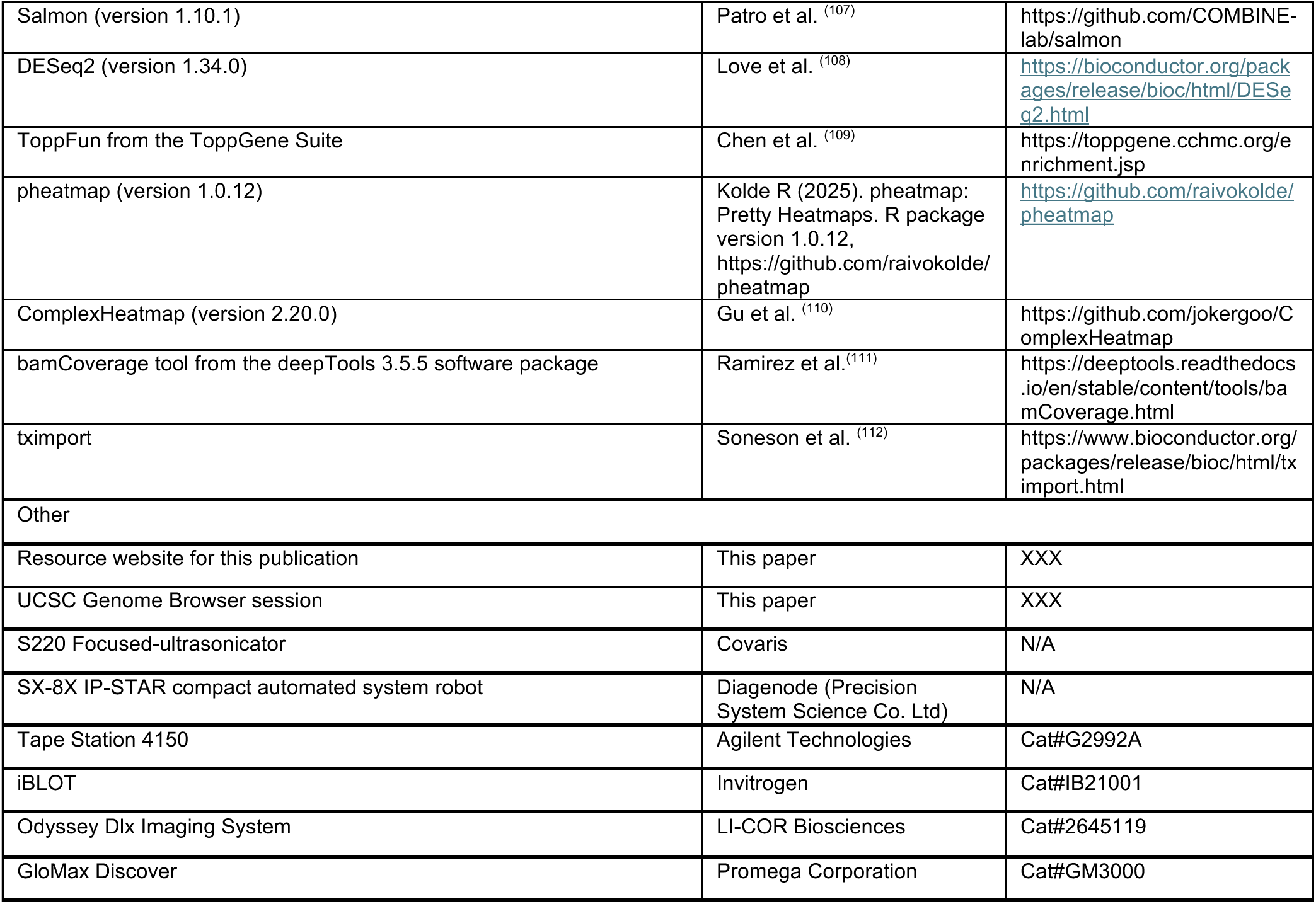

### EXPERIMENTAL MODEL AND STUDY PARTICIPANT DETAILS

Flp-In-293 and Flp-In T-REx-293 cells were maintained in DMEM supplemented with 10% FBS, 0.1% Anti-anti and 0.2% normocin. Similarly, the vTR stable cell lines containing the GFP fusions were grown in DMEM supplemented with 10% FBS, 0.1% Anti-anti, 0.2% normocin, and 200 µg/mL of hygromycin B. All cells were maintained in incubators at 37°C and 5% CO2. The cells were routinely tested and confirmed to be negative for mycoplasma contamination.

### METHOD DETAILS

#### vTR plasmid production

The pcDNA5/FRT/TO backbone and DNA sequences for EBV vTRs from strain B95-8 were supplied to Twist Bioscience to generate untagged, N-terminally GFP tagged and C-terminally GFP tagged plasmids. The coding sequence was codon optimized if necessary for production by Twist Bioscience. Some of the vTR constructs could not be generated by Twist Bioscience, so they were made through restriction digest or TOPO cloning into the same plasmid backbone instead (labeled in the resource table as “This paper”). We used AlphaFold3 (97) to determine whether the N-terminal or C-terminal fusion of GFP would have noticeable effects on the structure of each vTR protein. Plasmids utilized in this manuscript have been deposited in Addgene for distribution to other labs.

#### Transient transfection

Flp-In-293 cells were plated in 6 well plates at 500,000 cells/well in DMEM with 10% FBS and 0.1% Anti-anti. The following day cells were transfected (3 wells per vTR plasmid) with Lipofectamine 2000, according to the manufacturer’s protocol, using 0.8 mg of vTR plasmid and 0.2 µg of puromycin plasmid. The day after the transfection, each 6 well of cells was transferred to a well of a 12 well plate in media containing 2 µg/mL puromycin to kill off non-transfected cells. After 24 hours of selection, the media was changed to regular growth media and cells were allowed to recover for 24 hours before RNA collection using a mirVana RNA isolation kit, following the manufacturer’s protocol.

#### Stable cell generation

Flp-In-293 cells were plated on 6 well plates in DMEM supplemented with 10% FBS, 0.1% Anti-anti, and 0.2% normocin. Two days later, the cells were transfected using Lipofectamine 2000, according to the manufacturer’s protocol, with 0.25 µg of vTR plasmid and 2.25 mg of the pOG44 Flp-Recombinase expression vector. Within three days, the media was changed to media containing hygromycin at 50 mg/mL and the hygromycin was gradually increased to 200 mg/mL to produce a pool of cells stably expressing the vTR. The purity of the pool was confirmed by microscopy for GFP and Western blotting using an anti-GFP antibody to confirm expression of vTR fusions of the expected size. Doxycycline inducible Flp-In T-REx-293 cells were used to generate cells stably expressing EBNALP and were cultured in media also containing 100µg/mL zeocin. These cells were cultured in 1µg/mL doxycycline for overnight prior to collection.

#### Genomic assay sample collection

To ensure that the different vTR expressing cell lines were collected under similar culture conditions, stable cell lines were grown to confluency in four T175 flasks before transferring 90 million cells to six 15 cm dishes (15 million cells/dish). After incubating overnight, cells were trypsinized and two dishes/replicates were combined and standardized to 23 million cells per replicate for RNA-seq, ChIP-seq, and ATAC-seq experiments.

#### RNA-seq

Three million cells were washed with PBS and total RNA was isolated using the mirVana RNA extraction kit according to manufacturer’s instructions. RNA quality was measured using the 4150 TapeStation System and samples were sent to either the Cincinnati Children’s Hospital Medical Center (CCHMC) Genomics Sequencing Facility or Novogene for sequencing on either an Illumina NovaSeq 6000 or NovaSeq X Plus (paired-end, 150 bp read length).

#### ChIP-seq

20 million cells were resuspended in 20 mL of media for a final concentration of 1 million cells/mL. To crosslink the cells, 2 mL of 11% formaldehyde, 0.1M NaCl, 1 mM EDTA, 0.5 mM EGTA, and 50mM HEPES (pH 8.0) were added to the 20 mL of cells in media and crosslinked while rotating at room temperature for 10 minutes. The crosslinking reaction was stopped by adding 1.1 mL of 2.5M glycine and rotating for 5 minutes at room temperature. The cells were washed 2 times with cold PBS and then pelleted by centrifugation at 5000 RPM for 5 minutes and then stored in an Ultra-low freezer until further processing for ChIP. Crosslinked nuclei were prepared by washing the cell pellet with 1 mL of ChIP L1 buffer (10mM HEPES pH 8.0, 140 mM NaCl, 1 mM EDTA, 10% glycerol, 0.25% Triton X-100, and 0.5% NP-40) plus protease inhibitors and rotating at 4 °C for 10 minutes and pelleted by centrifuging at 10,000 g for 5 minutes. Then the cell pellets were washed with 1 mL of ChIP L2 buffer (10 mM Tris pH 8.0, 1 mM EDTA, 200 mM NaCl, and 0.5 mM EGTA) plus protease inhibitors by rotating at room temperature for 10 minutes and the nuclei were pelleted by spinning at 10,000 g for 5 minutes. The nuclei were then washed with 0.5 mL of sonication buffer (TE + 0.1% SDS) with protease inhibitors and again pelleted by centrifugation. The pelleted nuclei were resuspended in 1 mL of sonication buffer containing protease inhibitors and sonicated using a Covaris S220 focused-ultrasonicator at 10% duty cycle, 175 peak power, 200 burst/cycle for 7 minutes. The sonicated chromatin was centrifuged at maximum speed for 10 minutes and the supernatant of soluble chromatin was collected. The salt and detergents were adjusted to final concentrations of 150 mM NaCl, 0.1% NaDOC, 1% Triton X-100, 5% glycerol and protease inhibitors. The chromatin was pre-cleared of Ig by adding 20 µL of Dynabeads Protein A or G and rotating at 4 °C for 40 minutes and centrifuged for 10,000 g for 10 minutes. We performed the ChIP using 200 mL of the pre-cleared chromatin on a SX-8X IP-STAR compact automated system robot from Diagenode along with the antibody against the protein of interest and Dynabeads. For the vTRs, we used the Abcam anti-GFP antibody. For RBPJ we used the Cell Signaling Technologies RBPSUH (D10A4) XP antibody. The ChIP material was eluted using 10 mM TrisHCl and libraries were prepared by ChIPmentation (113) and sent to the CCHMC Genomics Sequencing Facility for sequencing on either an Illumina NovaSeq 6000 or NovaSeq X Plus (single-end, 100 bp read length).

#### ATAC-seq

We used 50,000 cells and followed the OMNI ATAC protocol (114) to generate the libraries of open chromatin that were subsequently sequenced. Briefly, transposase (Tn5) loaded with sequencing adapter sequences was used to cut the genome and insert sequencing adapters to tag the accessible DNA, as detailed in Buenrostro et al. (115). The resulting accessible DNA sequences were isolated, and libraries were prepared using the OMNI ATAC protocol (114). The libraries were sent to the CCHMC Genomics Sequencing Facility for sequencing on an Illumina NovaSeq X Plus (paired-end, 150 bp read length).

#### Western blots

Cells were lysed in RIPA-DOC-PPI and sonicated, and protein concentration was determined using a BCA protein assay. 10 μg of lysates were loaded in 4-12% NuPAGE Bis-Tris gels, run in MOPS buffer, and transferred to nitrocellulose using an iBlot. The membranes were blocked for 1 hour in Intercept Blocking Buffer and incubated overnight at 4°C in Intercept Blocking Buffer + 0.1% Tween 20 with anti-GFP or actin antibodies diluted to 1:1000. The next day, the membranes were washed 2-3 times for 10 minutes with PBS + 0.1% Tween 20. The washed membranes were then incubated with rotation for 45 minutes with fluorescent secondary antibodies (diluted to 1:10,000 in Intercept Blocking Buffer + 0.1% Tween + 0.01% SDS). After three more 10 minute washes in PBS + 0.1 Tween 20, the membranes were imaged using an Odyssey DLx Imaging System.

#### Co-immunoprecipitation assays

10 million Flp-In-293 parental cells (control) or RTA-GFP expressing cells were lysed in 1 mL of co-immunoprecipitation buffer (25 mM HEPES, pH 7; 200 mM KCl; 20 mM NaF; 0.2mM EGTA; 1% NP40). The lysates were sonicated in a water bath sonicator for 4 minutes of 10 seconds on/off cycles and then centrifuged for 5 minutes at 12,000 RPM to pellet debris. We combined 100 µL of the soluble lysate with 100 µL of 2X Laemmli sample buffer to use as inputs and then used 500 µL of the soluble lysates for the co-immunoprecipitation through the addition of 25 µL of mouse anti-GFP antibody conjugated to magnetic beads and incubated rotating at 4 degrees for 2 hours. The beads were isolated with a magnet and washed 5 times with 500 µL of co-immunoprecipitation buffer. The immunoprecipitated material was eluted off the beads with 50 µL of 2X Laemmli sample buffer. We then loaded 10 µL of inputs and IP elutions on 4-12% NuPAGE Bis-Tris gradient gels, transferred to nitrocellulose and probed with rabbit anti-GFP or rabbit anti-RBPJ antibodies. The blots were washed 3x for 10 minutes, incubated with HRP conjugated anti-rabbit secondary antibodies, washed again and then imaged using enhanced chemiluminescent HRP substrate.

#### Split NanoLuc luciferase protein complementation assays

For the split NanoLuc luciferase experiments, 200,000 Flp-In-293 cells were plated per well of a 12 well plate. The next day, the cells were transfected with 0.2 mg/well of the indicated split NanoLuc luciferase constructs (LgBit fused to the various RTA constructs and SmBit fused to RBPJ). The next day the wells were trypsinized and 50,000 cells were plated/well of a 96 well in quadruplicate wells. After allowing the cells to attach for 24 hours, the media was changed to 50 mL of Opti-Mem + 12.5 mL of Promega Nano-Glo Live Cell reagents. The plate was then read over a 10 second integration on a GloMax (Promega) plate reader. The signals from all the wells were normalized to the average of the cells transfected with only the RTA-LgBit fusion to control for background signal.

### QUANTIFICATION AND STATISTICAL ANALYSIS

#### RNA-seq data analysis

RNA sequencing data were processed using the nf-core/rnaseq pipeline (version 3.11.1) https://doi.org/10.5281/zenodo.7789554). Initial quality control of the raw sequencing data was performed using FastQC (version 0.11.9) (https://www.bioinformatics.babraham.ac.uk/projects/fastqc/). Low-quality bases and adapter sequences were then trimmed and filtered from the reads using Cutadapt (version 3.4) (https://doi.org/10.14806/ej.17.1.200) and Trim Galore (version 0.6.7) (https://doi.org/10.5281/zenodo.5127899), respectively. Ribosomal RNA sequences were subsequently removed from the aligned data using SortMeRNA (version 4.3.4) (100) against the default libraries augmented with human rRNA sequences to eliminate potential contamination from non-target RNA species. STAR (version 2.7.9a) (101) was used to align the trimmed reads to the GRCh38 genome (https://ftp.ebi.ac.uk/pub/databases/gencode/Gencode_human/release_40/GRCh38.pri mary_assembly.genome.fa.gz) with GENCODE release 40 annotations (https://ftp.ebi.ac.uk/pub/databases/gencode/Gencode_human/release_40/gencode.v40.annotation.gtf.gz) (116). The resulting alignments were sorted and indexed using SAMtools (version 1.16.1) (102). To further evaluate the quality of the RNA sequencing data, several tools were employed, including RSeQC (version 3.0.1) (103), Qualimap 2 (version 2.2.2-dev) (104), dupRadar (version 1.28.0) (105), and preseq (version 3.1.1) (106). These tools provide comprehensive assessments of various quality metrics, such as read distribution, GC content, duplication rates, and library complexity, ensuring reliable data for downstream analysis. For transcript quantification, Salmon (version 1.10.1) (107) was used to estimate the expression levels of transcripts. Differential gene expression analysis was performed using DESeq2 (version 1.34.0) (108). Genes were considered differentially expressed if they had a 20% fold change compared to the GFP control and an adjusted p-value threshold of less than 0.05. The EBNALP stable cell line was compared to the Flp-In T-REx-293 GFP control cell line. The other stable cell lines were compared to the Flp-In-293 GFP control cell line. For the transient transfections, the untagged vTR was compared to the transient GFP control. Transcript abundance estimates were generated by Salmon and prepared for DESeq2 analysis using the tximport package (112). Gene-level summarization of transcript quantifications was performed during import using annotation information provided by the GENCODE GTF file, which contained 61,544 entries. Pathway enrichment analyses for the differentially expressed genes were performed using ToppFun from the ToppGene Suite (109) and the GO Consortium’s Generic GO subset (goslim_generic) terms annotated as a biological process ((117, 118) and https://doi.org/10.5281/zenodo.14083199). The parameters used for ToppFun were “Probability density function” for the pValue Method, “GO: Biological Process for the Feature”, “None” for the Correction, “1.0” for the p-Value cutoff, and gene limits of 5 and 2,000. The ToppFun calculated background set of 20,262 genes was used. Each set of DEGs for a specific vTR and direction (e.g. BMRF1 up-regulated genes) was processed separately, using the Ensembl IDs as the input set. The results were filtered against the GO slim terms and an adjusted p-value was calculated using the Benjamini-Hochberg Procedure. The odds ratio was calculated using the formula from Enrichr (117, 118), specifically (1.0 * a * d) / max(1.0 * b * c, 1), where a is “the number of overlapping genes”; b is “the number of genes in the annotated set” - “the number of overlapping genes”; c is “the number of genes in the input set” - “the number of overlapping genes”; and d is “the number of genes in the background set” - “the number of genes in the annotated set” - “the number of genes in the input set” + “the number of overlapping genes”. An adjusted p-value threshold of 0.05 was used to generate the pathways enrichment figures. Heatmaps were made with pheatmap (version 1.0.12) (119), except for **Figure 7B**, which was made with ComplexHeatmap (version 2.20.0) (110).

#### Data Visualization

Visualization tracks for each dataset were created for the UCSC Genome Browser (94). Signal tracks (in bigWig format) were created using the bamCoverage tool from the deepTools 3.5.5 software package (111) with the parameters --normalizeUsing CPM and --binSize 10. For the RNA-seq signal tracks, the command was run twice in order to maintain strandedness. For the forward strand track, the parameter --filterRNAstrand forward was included. For the reverse strand track, the parameters --filterRNAstrand reverse and --scaleFactor -1 were included. Genome browser screenshots were made with the Integrative Genomics Viewer (120).

#### Chromatin State Analysis

Regions corresponding to particular chromatin states in HEK-293 cells were derived previously (61). Briefly, a ChromHMM model was trained with the histone marks H3K4me1, H3K4me3, H3K36me3, H3K27ac, H3K9me3, and H3K27me3, CTCF ChIP-seq, ATAC-seq, and GENCODE promoter regions. ChIP- and ATAC-seq peaks from our analyses were assigned to a ChromHMM state using “bedtools intersect” (version 2.31.1) (121), and total coverage was derived by using “bedtools groupby” to group intersected regions by their label and sum the sizes of intersecting peaks. Chromatin state distribution over accessible chromatin was calculated in the same way, using the ATAC-seq peaks from the monoGFP (control) experiment. The total size of each chromatin state across the genome was assessed similarly by first removing the ENCODE blacklist regions from the ChromHMM states using “bedtools subtract” and then aggregating with “bedtools groupby”.

#### Transcription factor motif enrichment analyses

A modified version of HOMER v4.9.1 (122) which utilizes a log2 scoring system was used for all motif enrichment analyses. Known motif enrichment using CisBP v3.00 motifs (123) was computed for each peakset with findMotifsGenome.pl and the “-size given” parameter. The enrichment heatmaps in **Figures 2F** and **4** were constructed by first identifying the top (lowest p-value) 10 motifs in each peakset that come from distinct motif families. The Pearson correlation of the normalized motif p-values for each motif was calculated, and the motif with the highest correlation was iteratively removed until the highest correlation coefficient was below 0.85. *De novo* motif enrichment of RTA was performed with the HOMER “-size given” parameter and the CisBP 3.0 motifs in the “-mcheck” parameter.

#### ChIP-seq peak calling

ChIP-seq alignment and peak calling was performed using a version of the ENCODE ChIP-seq pipeline (57, 58, 123) modified to include HOMER-based CIS-BP motif enrichment and to run within a Nextflow environment. All alignments were performed to the hg38 genome. Each vTR used the GFP-expressing cell line as a background control for peak calling. For all downstream analyses, “IDR conservative” peaks were used.

#### ATAC-seq peak calling and differential analysis

ATAC-seq alignment and peak calling was performed using a version of the ENCODE ATAC-seq pipeline (57, 58, 123) slightly modified to run within a Nextflow environment. All alignments were performed to the hg38 genome. For all downstream analyses, “IDR conservative” peaks were used.

Differentially accessible regions were identified using DiffBind v3.12.0 (124) with an FDR threshold of 0.05. Assignments to “direct” and “indirect” chromatin effects were made using bedtools v2.31.1 (121) and command lines similar to “bedtools intersect -a $open_bed -b $chip_peaks -wa -u > $vtr_direct_opening.bed” or “bedtools intersect -a $open_bed -b $chip_peaks -v > $vtr_indirect_opening.bed”.

#### BMRF1 disease associations

The Regulatory Element Locus intersection (RELI) algorithm (16) was used to compare BMRF1 ChIP-seq peaks with tag SNPs for phenotypes in the GWAS Catalog (125) (downloaded on February 18, 2025). The final set of 439 tested phenotypes consisted of those that 1) describe disease processes, 2) have at least 10 tag SNPs which are not in linkage disequilibrium (r^2^ < 0.2), and 3) were assessed in populations of primarily European ancestry (to match the ancestral background of our cell line).

The BMRF1 interaction network (**Figure 6D**) was constructed using STRING (126), with expert curation to assign group function. Variants that were overlapped by BMRF1 ChIP-seq peaks and were tag SNPs for one of the five rheumatoid arthritis-related phenotypes in **Figure 6C** were assessed for eQTL function using release 7 of the eQTL Catalogue (98). Gene targets of these eQTLs (p-value < 1E-5) were used as input to STRING.

## ADDITIONAL RESOURCES

Project website: XXX

UCSC Genome Browser session: XXX

