## Supplementary figures and images for "Systematic investigation reveals extensive Epstein-Barr virus transcriptional regulation of the human genome"

### Supplemental Figure 1

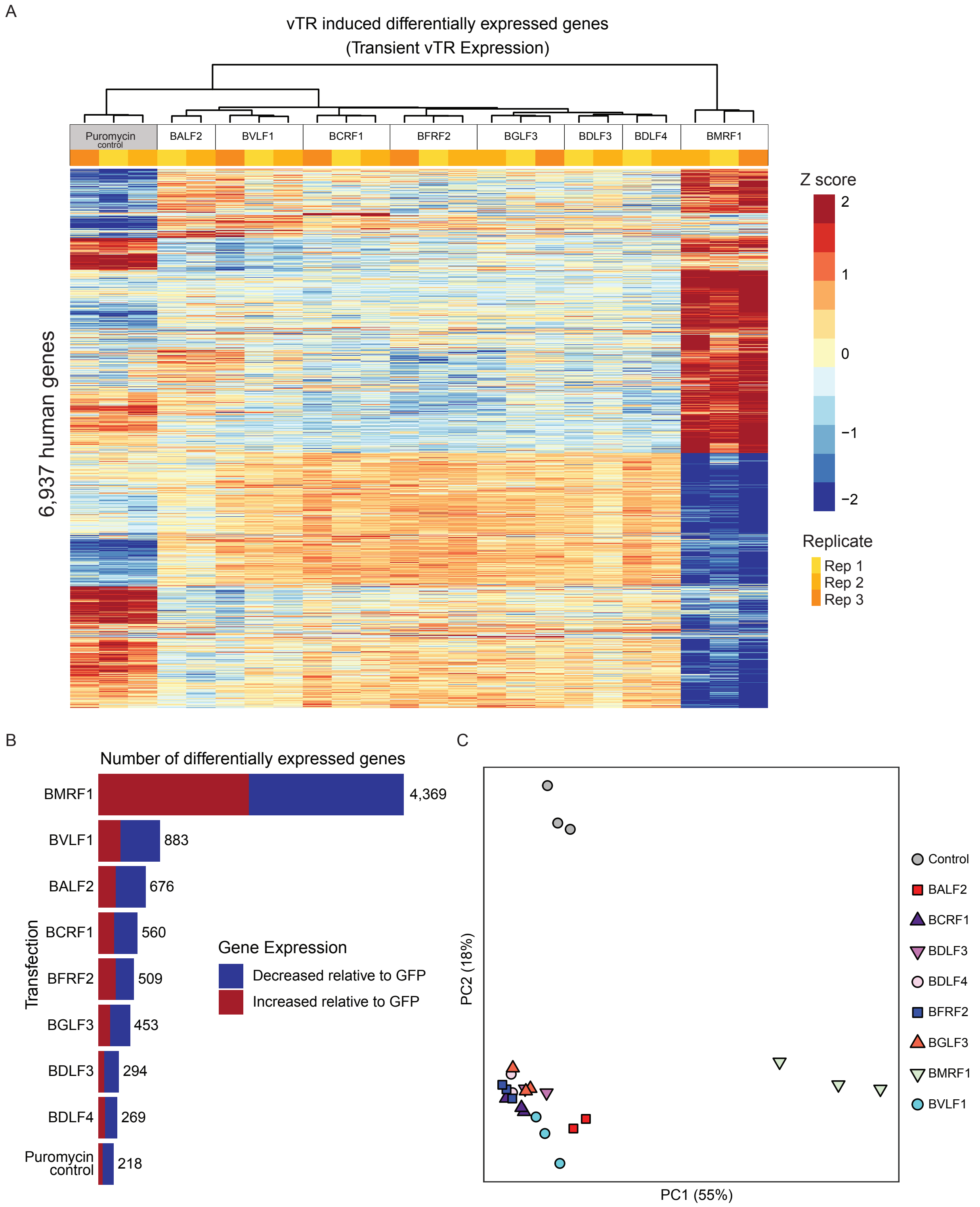

### Supplemental Figure 2

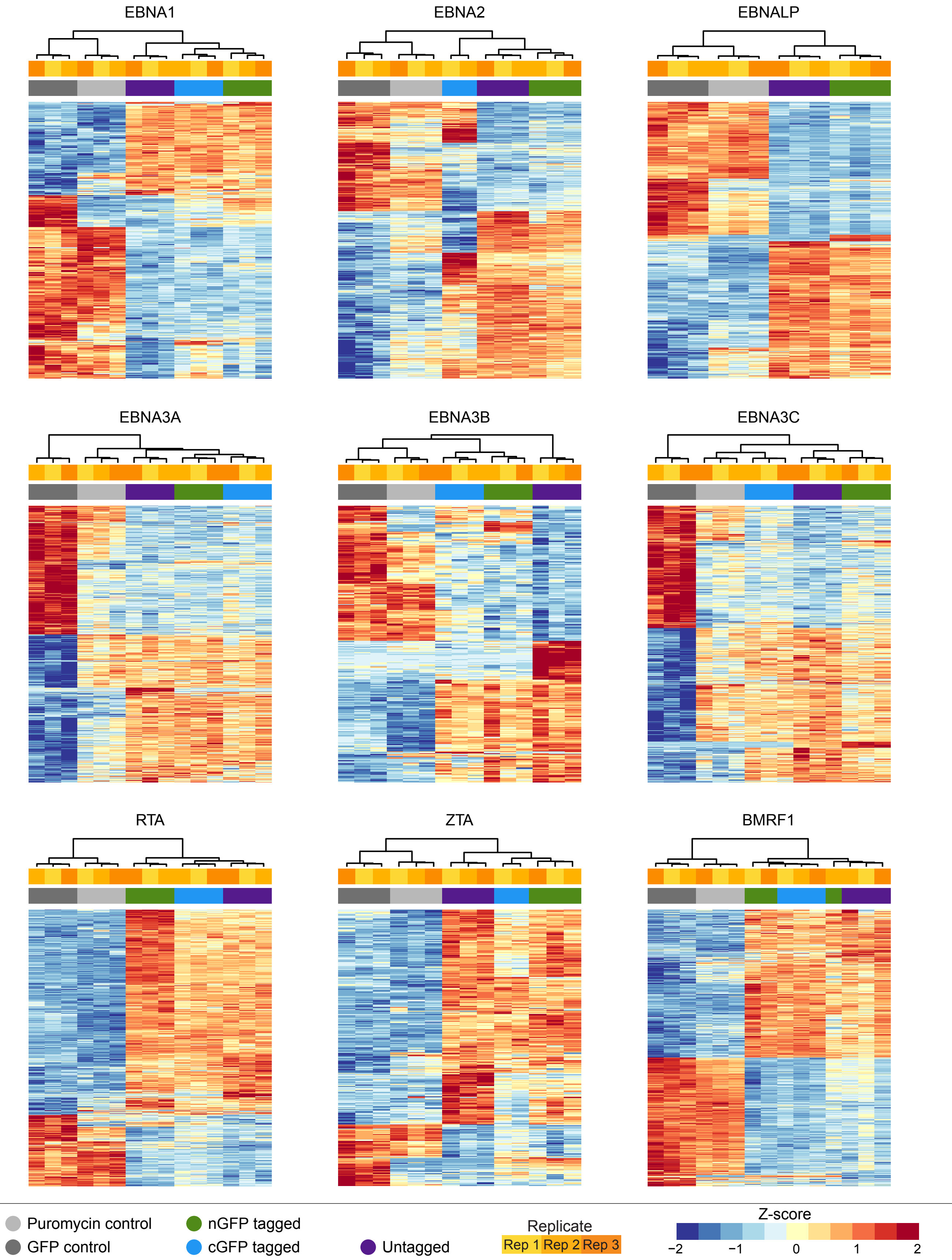

### Supplemental Figure 3

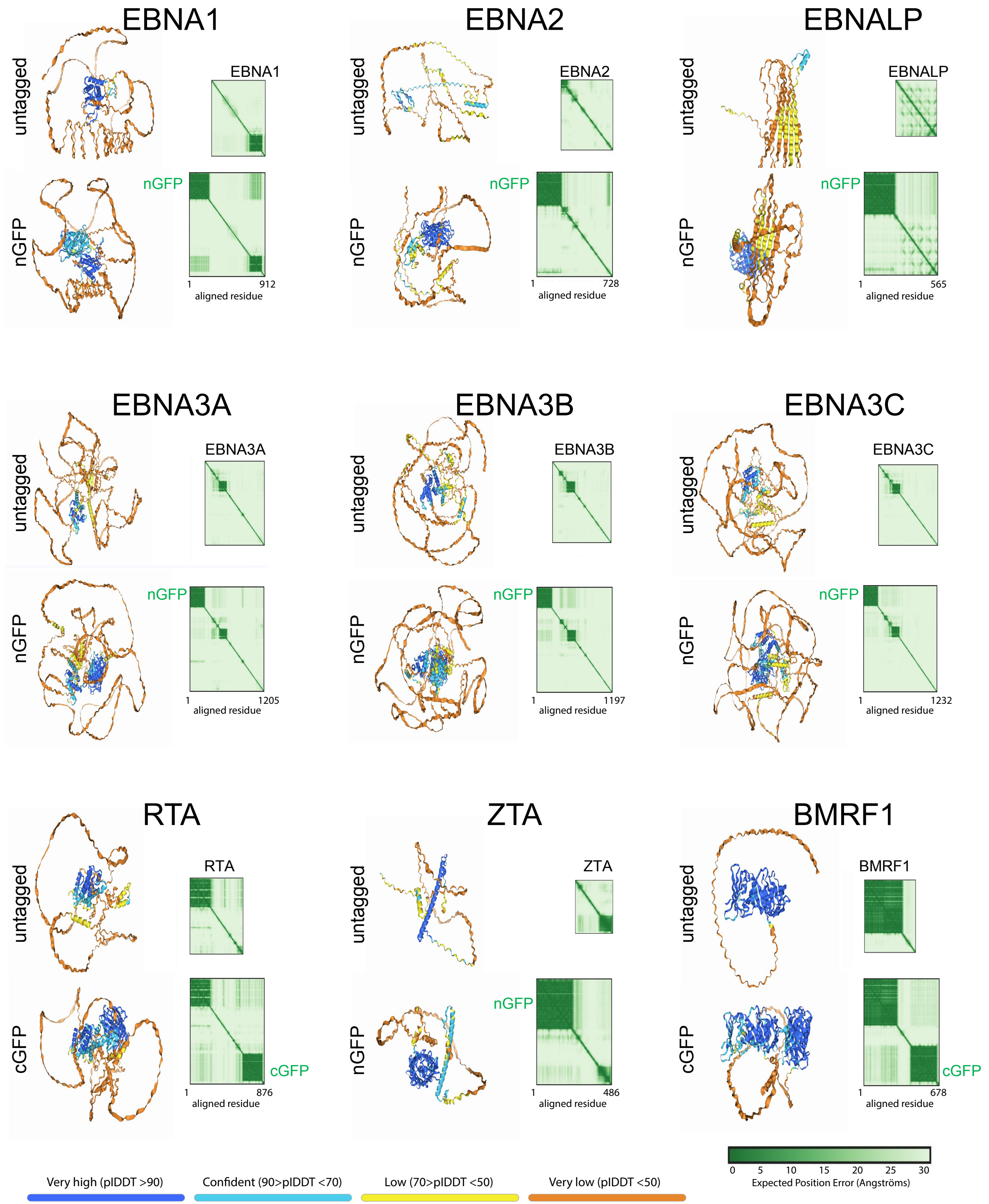

### Supplemental Figure 4

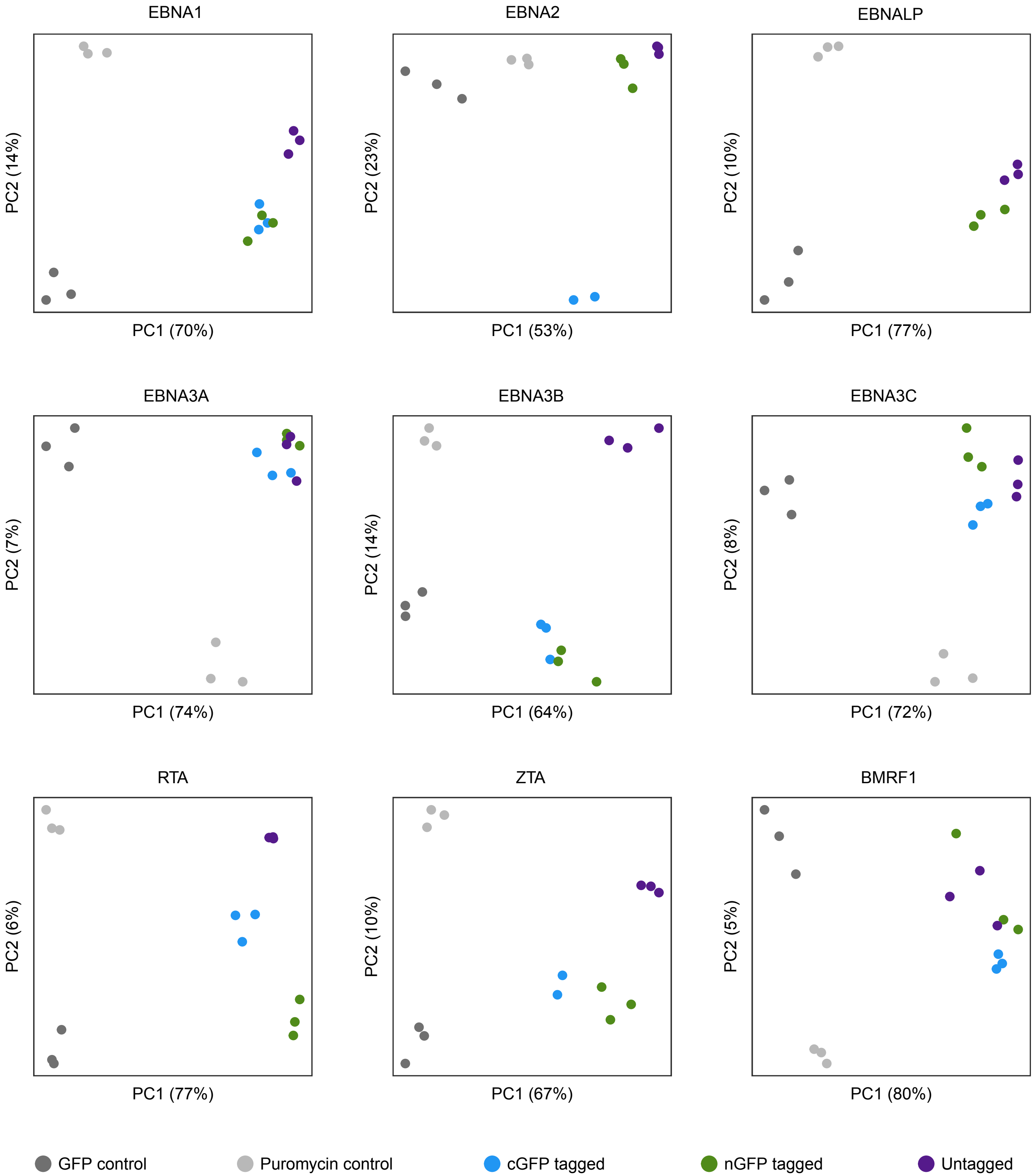

### Supplemental Figure 5

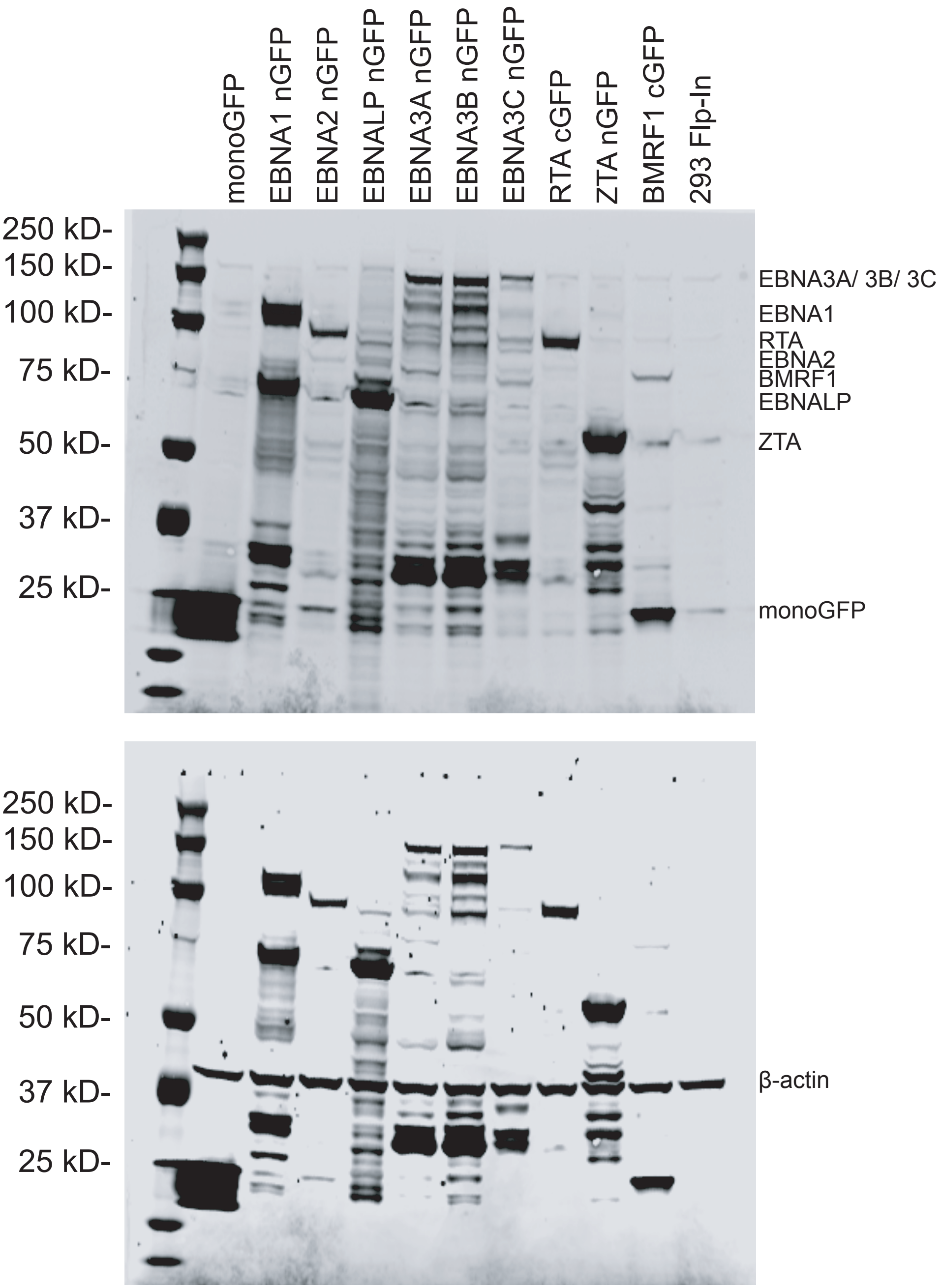

### Supplemental Figure 6

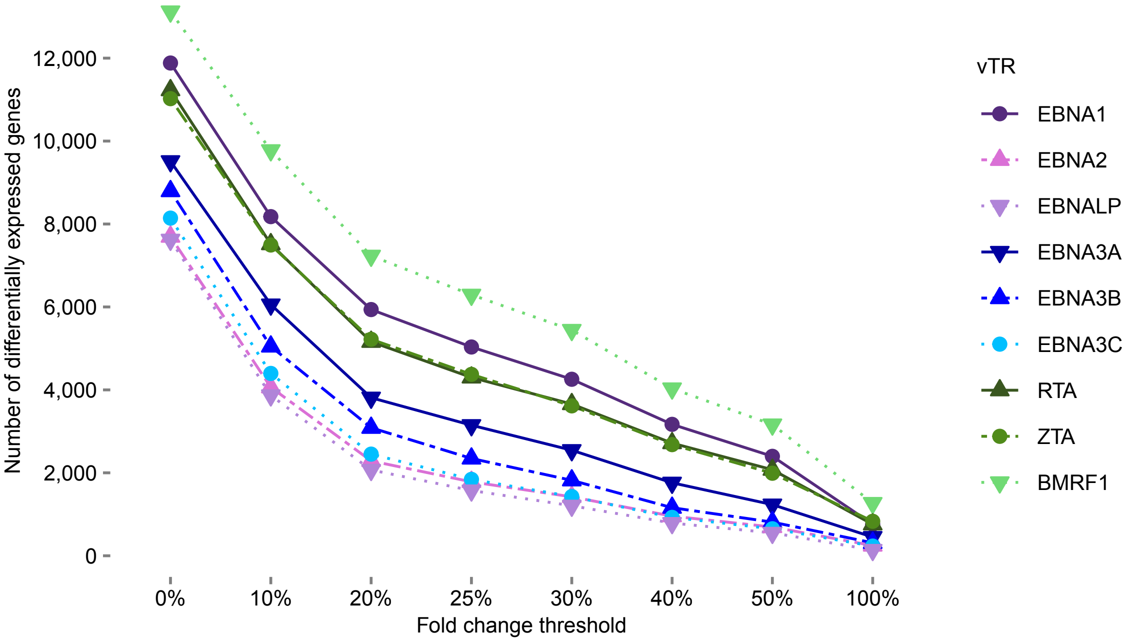

### Supplemental Figure 7

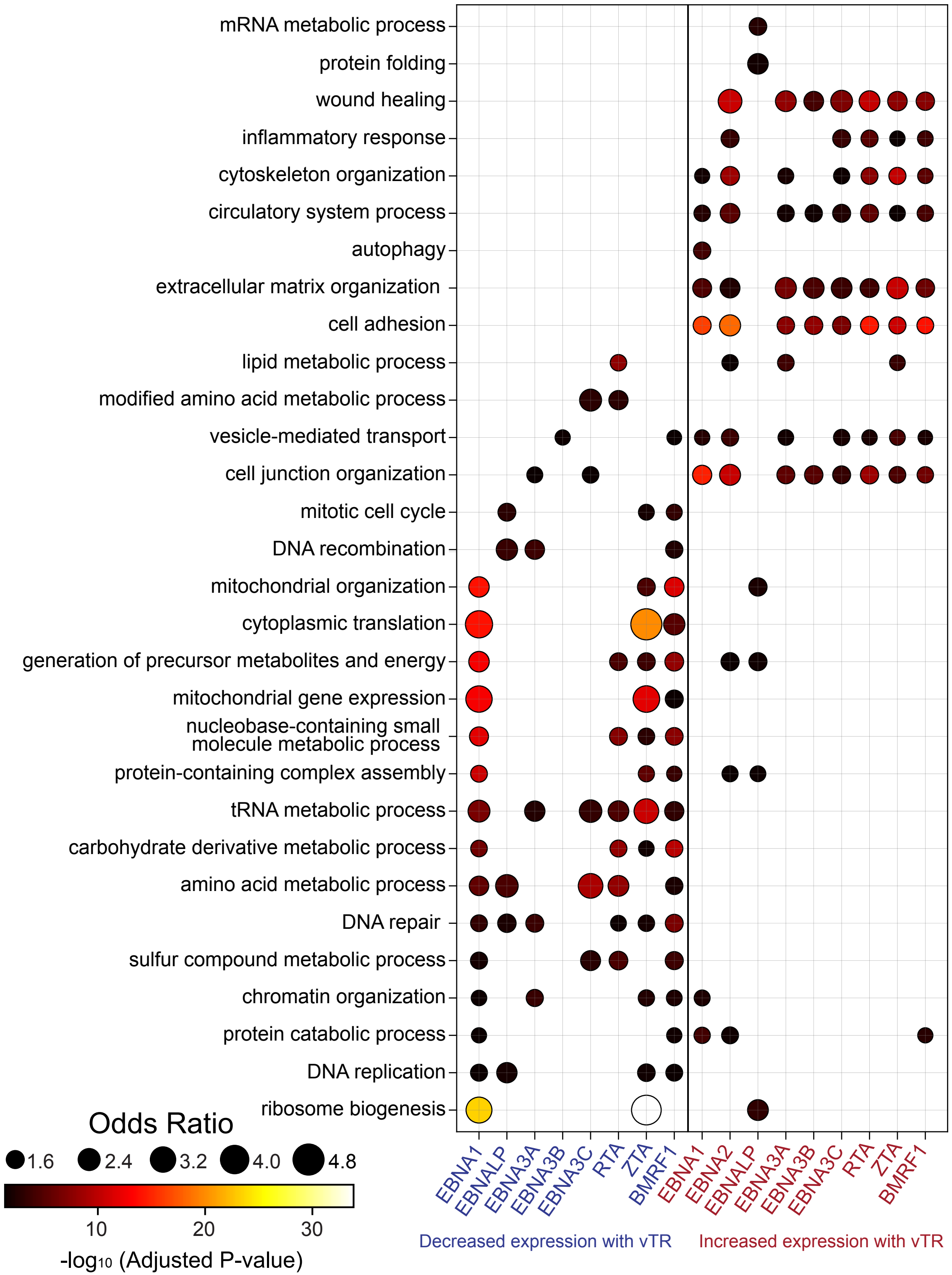

### Supplemental Figure 8

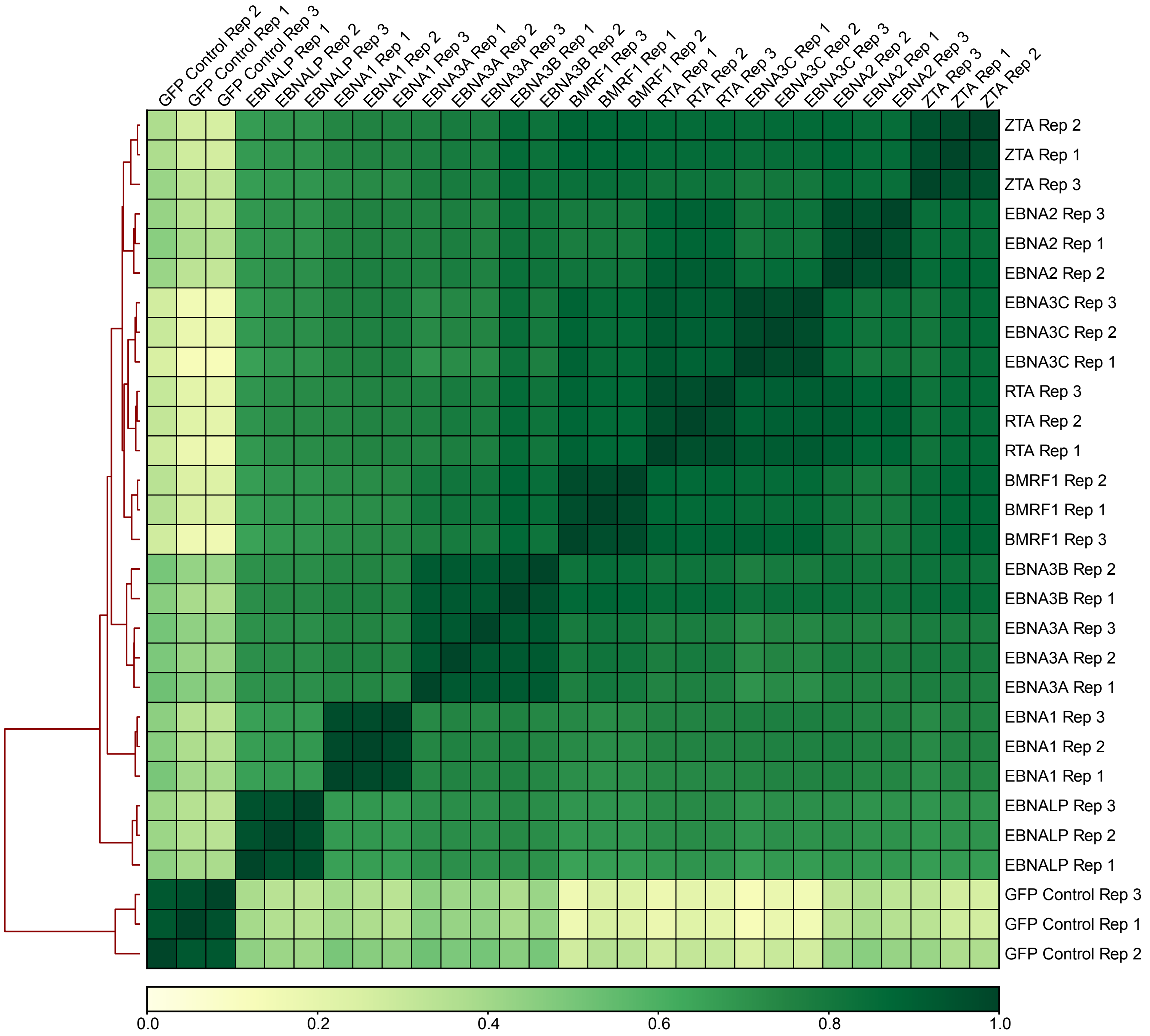

### Supplemental Figure 9

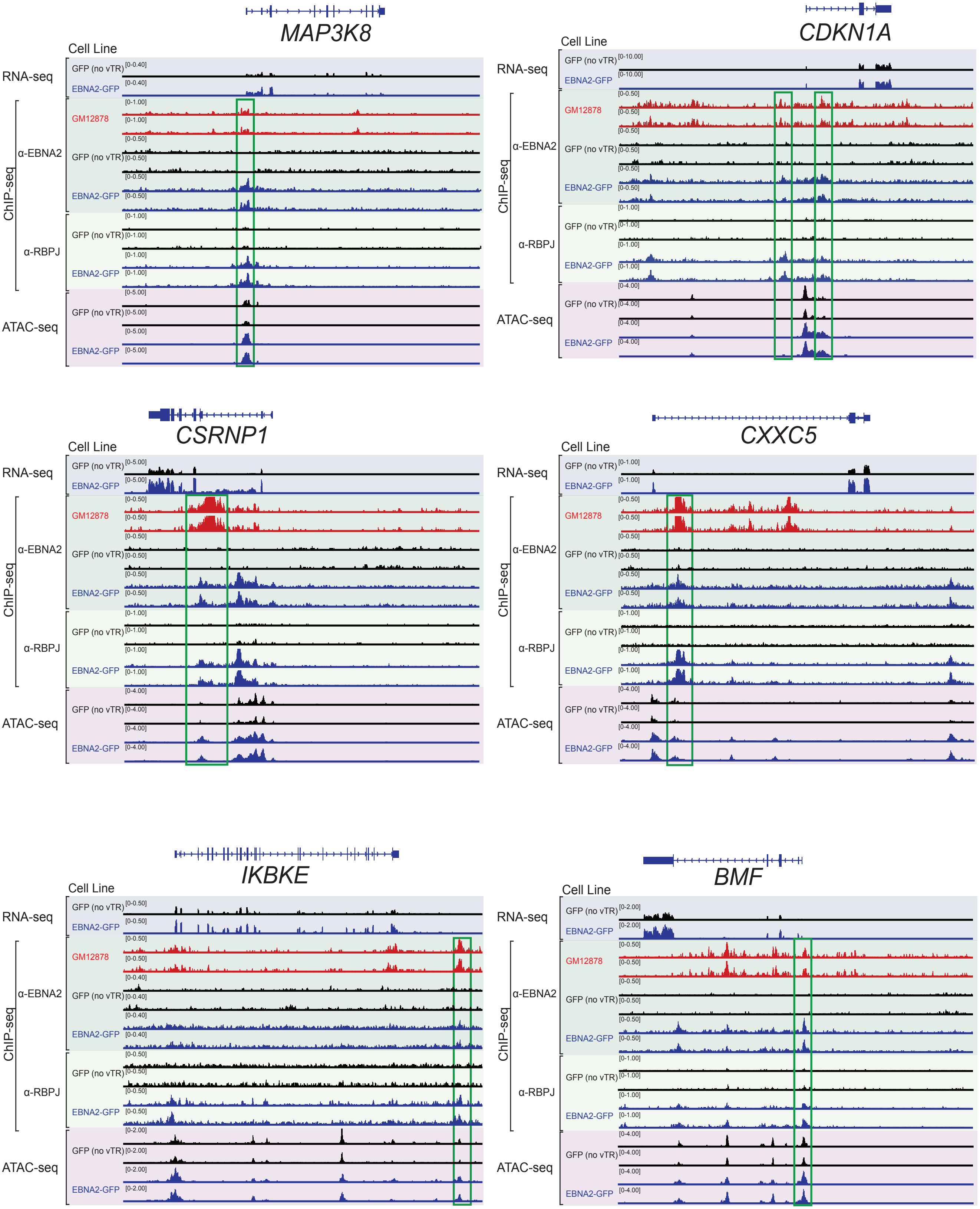

### Supplemental Figure 11

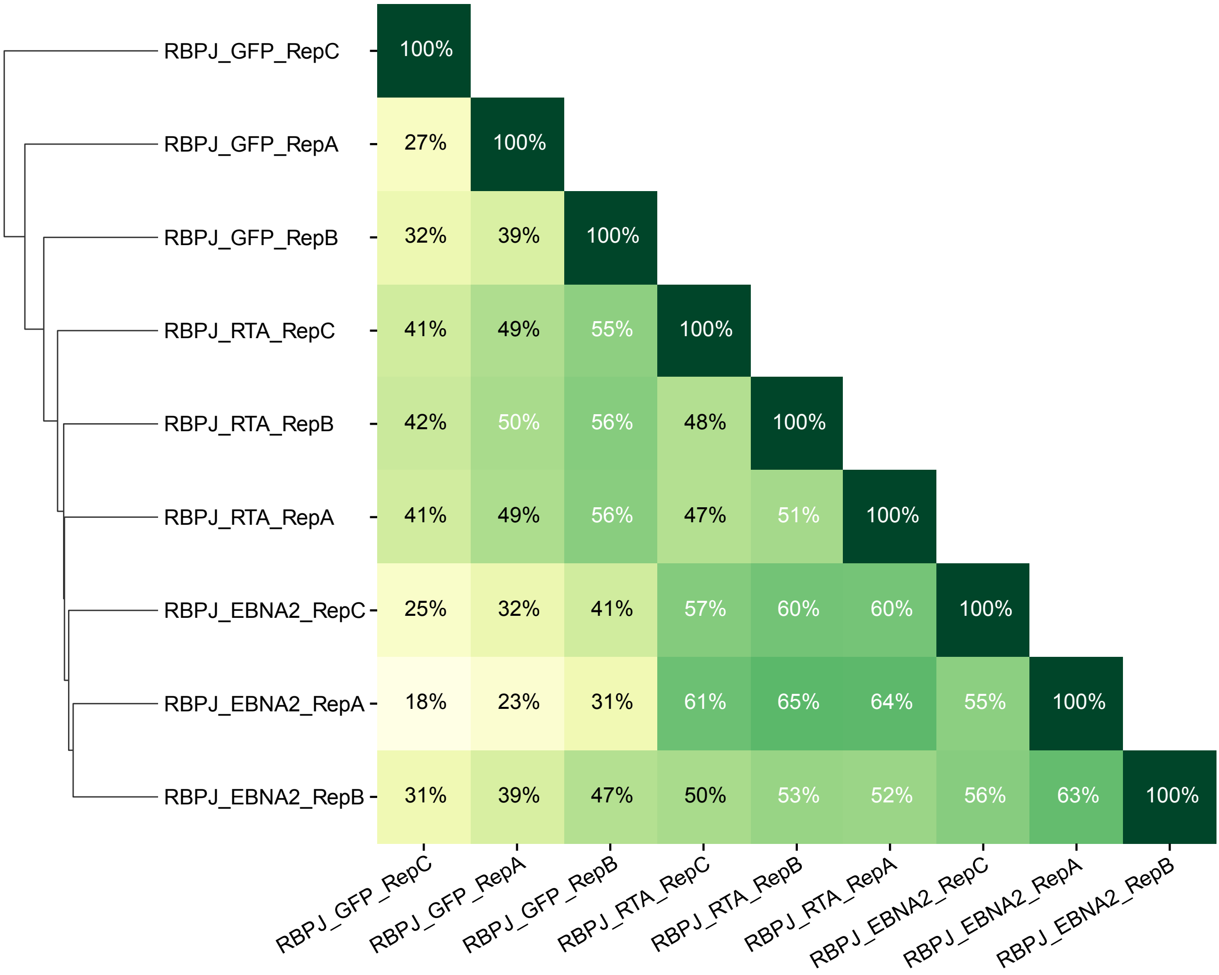

### Supplemental Figure 12

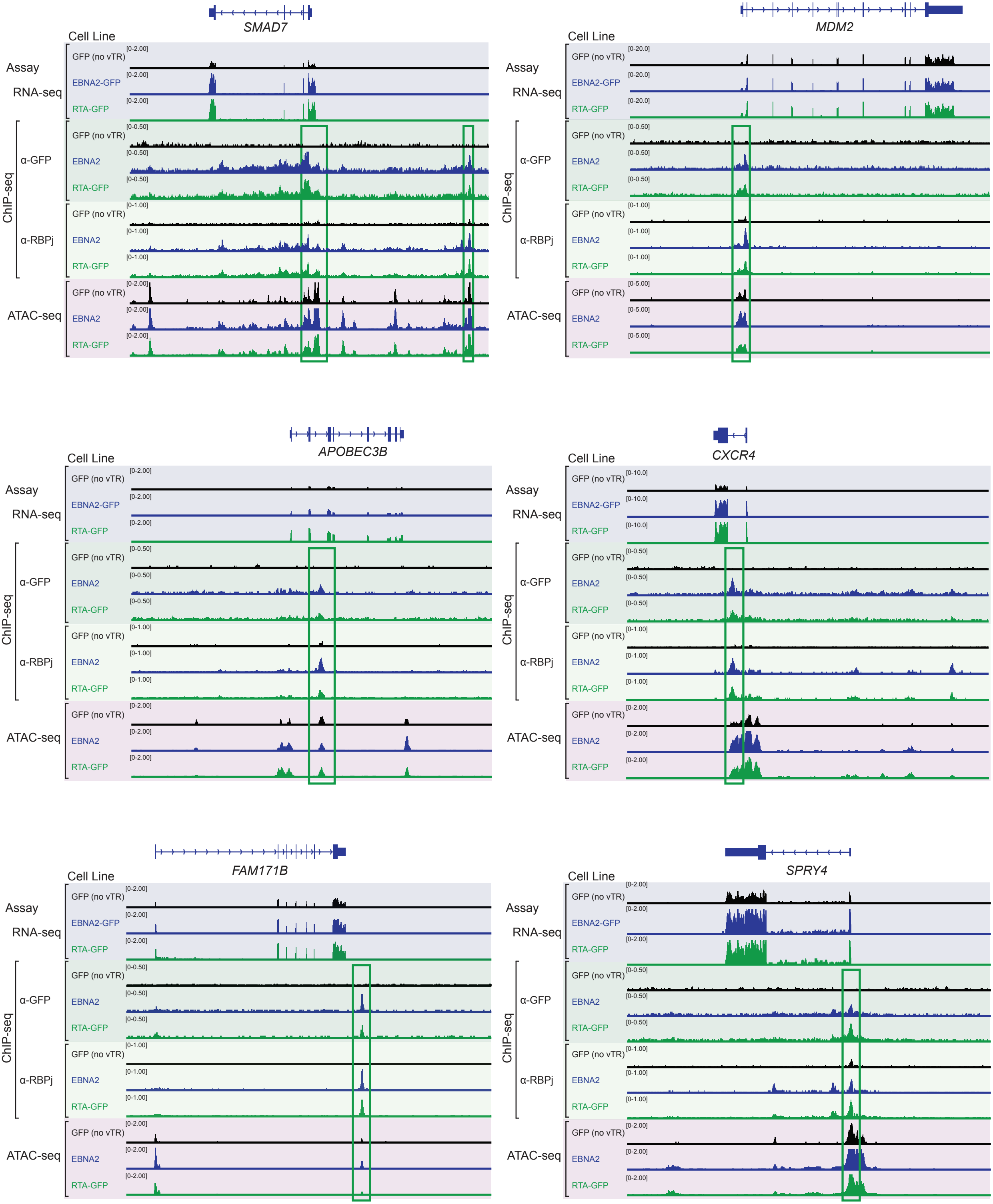

### Supplemental Figure 13

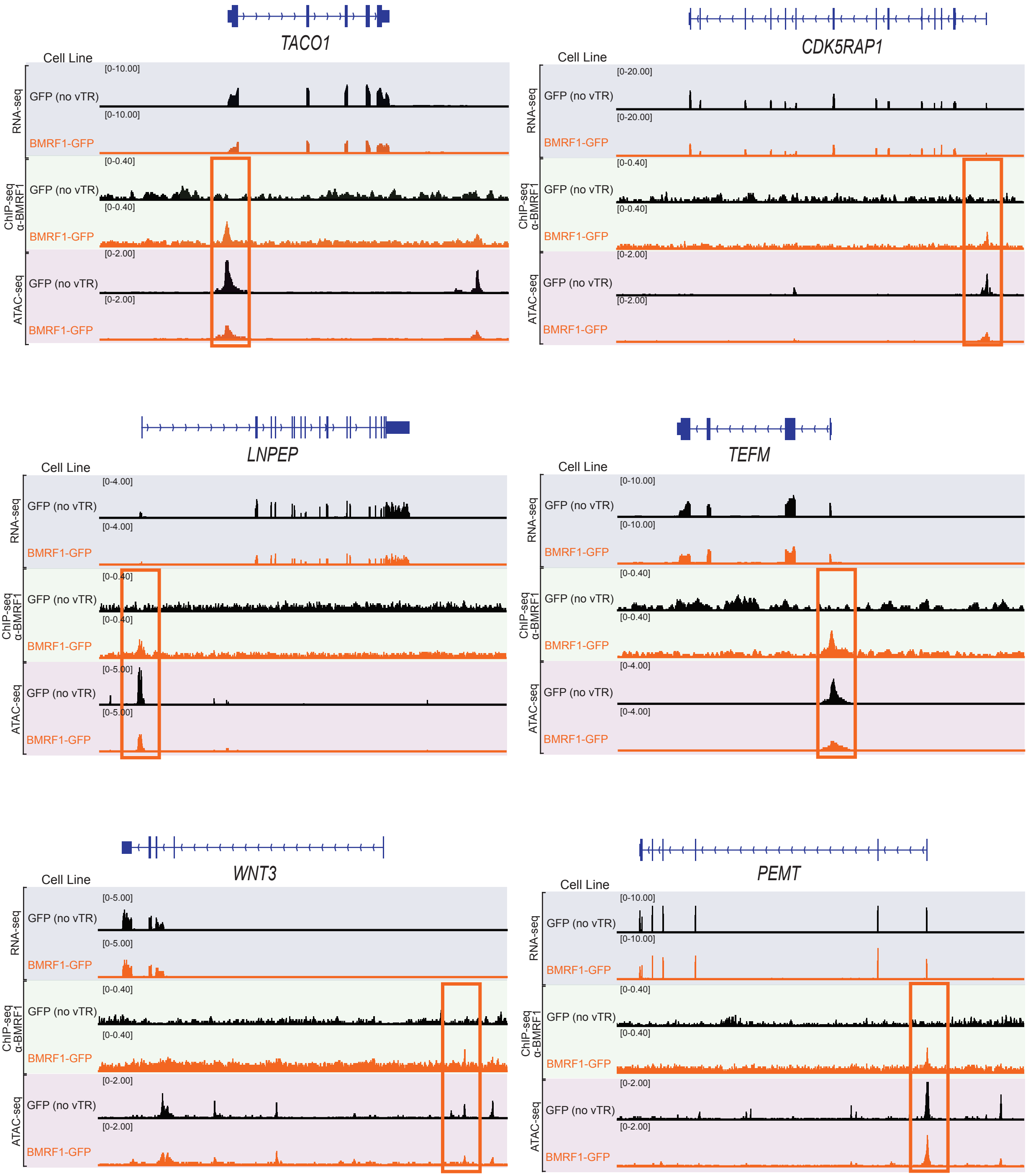
