## Supplemental Figure 10 for "Systematic investigation reveals extensive Epstein-Barr virus transcriptional regulation of the human genome"

A. Gene expression (RNA-seq) and genome binding (ChIP-seq)

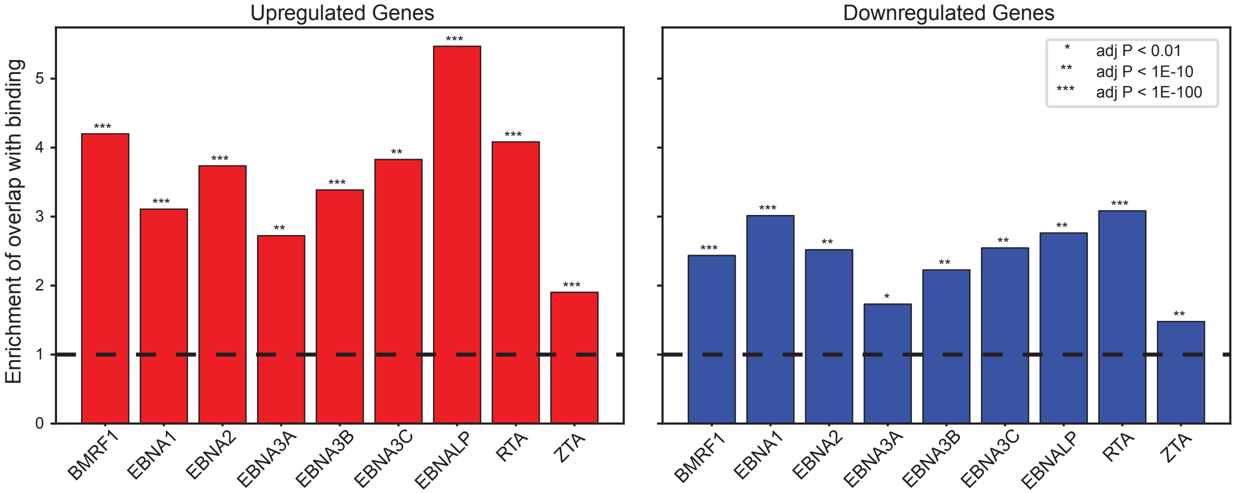


B. Chromatin accessibility (ATAC-seq) and genome binding (ChIP-seq)

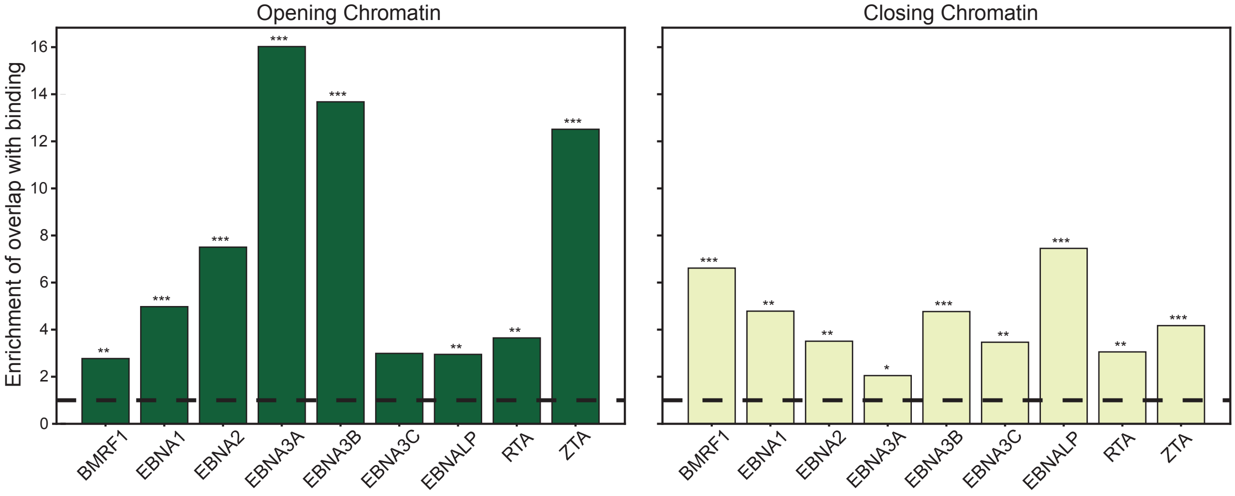
